# DLC1 loss drives multicellular streaming invasion by enforcing spatially coordinated Rho and β1 integrin signaling

**DOI:** 10.64898/2026.06.22.733747

**Authors:** Fiona Kühnel, Saskia Ebert, Oliver Piechota, Lisa Tellier, Florian Meyer, André Koch, Cristiana Lungu, Monilola A. Olayioye

## Abstract

Efficient cancer cell invasion requires coordinated control of actomyosin contractility, extracellular matrix (ECM) engagement and remodeling, yet how these processes are integrated in complex, three-dimensional (3D) environments remains unclear. Here, we identify the tumor suppressor and RhoGAP protein DLC1 as molecular brake on multicellular streaming invasion in collagen-rich ECM. Using CRISPRoff-engineered breast cancer spheroids, DLC1 reconstitution models and patient-derived organoids embedded in collagen gels, we show that DLC1 downregulation promotes an efficient multicellular streaming phenotype. This invasion program requires matrix metalloproteinase activity, β1 integrin engagement and Rho–ROCK-dependent actomyosin contractility. Mechanistically, DLC1 downregulation stabilized the rear-polarization of RhoA activity and increased β1 integrin abundance, plasma membrane localization and activation. Separation-of-function mutants revealed that DLC1 restrains invasion through a dual mechanism: its RhoGAP activity limited actomyosin-driven streaming, whereas its LD-like talin-binding motif controlled β1 integrin enrichment at the plasma membrane. Together, our findings provide a molecular basis for the prominent role of DLC1 as a metastasis suppressor.

## Introduction

Metastatic dissemination requires cancer cells to invade through three-dimensional (3D) extracellular matrices (ECM), where physical confinement and matrix architecture impose major constraints on cell migration (Paul et al. 2017; Yamada and Sixt 2019). In solid tumors, such as breast carcinomas, disease progression is accompanied by extensive remodeling of the stromal ECM, including increased deposition, crosslinking and alignment of fibrillar type I collagen. These changes alter the ECM mechanics and generate structural guidance cues that promote cell movement (Joshi et al. 2026; Bourgot et al. 2020). Within these environments, cancer cells adopt diverse invasion modes, including dissemination as single cells, collective groups, or multicellular streams, depending on the integration of cytoskeletal contractility, cell–matrix adhesion and proteolytic matrix remodeling (Talkenberger et al. 2017; Friedl and Gilmour 2009; Stehbens et al. 2024).

Cells sense and respond to matrix architecture and mechanics via integrins, transmembrane αβ heterodimers that link the ECM with the intracellular actin cytoskeleton via a set of adaptor proteins at so-called focal adhesions (Kanchanawong and Calderwood 2023; Saraswathibhatla et al. 2023). As the central β subunit of collagen receptors, β1 integrin plays a key role in adhesion signaling and traction-dependent cell migration (Chastney et al. 2025), and aberrant β1 integrin expression and activation have been linked to metastasis (Yao et al. 2007; Chastney et al. 2025).

Integrin function is controlled by receptor abundance and conformational activation, with clustering at sites of matrix engagement enhancing adhesion signaling and force transmission (Kanchanawong and Calderwood 2023). These processes are regulated through focal adhesion adaptor proteins such as talin and vinculin, and are coupled to downstream activation of Rho GTPase signaling that coordinates actin remodeling (Schwartz and Shattil 2000; Legerstee and Houtsmuller 2021). In particular, RhoA-mediated actomyosin contractility drives focal adhesion maturation and traction-dependent cell movement (Petrie and Yamada 2016). Aberrant activation of Rho signaling networks is commonly observed in cancer (Hodge and Ridley 2016), but how this feeds back to control β1 function during tumor cell invasion is incompletely understood.

Deleted in Liver Cancer 1 (DLC1), a RhoA-directed GTPase activating protein (GAP), has been established as a tumor suppressor in many cancer entities, including those of the breast (Xue et al. 2008; Wang et al. 2014). DLC1 localizes to focal adhesions through interactions with talin and tensin family proteins, thereby being at a critical interface between integrin signaling and the actin cytoskeleton (Li et al. 2011; Zacharchenko et al. 2016; Haining et al. 2018). While DLC1 loss has been associated with enhanced cell motility *in vitro* and metastasis *in vivo* (Holeiter et al. 2008; Goodison et al. 2005; Wang et al. 2014), the mechanisms by which DLC1 coordinates adhesion with contractility to control cell invasion programs in a 3D matrix environment have not been explored.

Here, we identify DLC1 as a central regulator of β1 integrin-dependent invasion in 3D. Using CRISPRoff-mediated epigenetic repression in MDA-MB-231 spheroids, DLC1 re-expression in bone-tropic MDA-MB-231 derivatives, and patient-derived breast cancer organoids, we show that DLC1 downregulation promotes multicellular streaming in fibrillar collagen. This is accompanied by enhanced collagen cleavage, stabilized polarization of RhoA activity and increased β1 integrin abundance, cell-surface localization and activation. Mechanistically, we find that DLC1 controls 3D cell invasion through a dual mode of action involving its GAP domain, which restricts spatial RhoA-dependent actomyosin contractility and its LD-like talin interaction motif, which regulates β1 integrin function. Together, our findings provide a molecular basis for the prominent role of DLC1 as a metastasis suppressor.

## Results

### DLC1 depletion promotes multicellular streaming and MMP-dependent invasion in collagen gels

Downregulation of DLC1 has been linked to metastasis (Goodison et al. 2005; Wang et al. 2014). However, the mechanisms through which DLC1 loss enables cancer cells to invade complex 3D ECM environments remain unclear. We addressed this using the highly invasive, triple-negative breast cancer (TNBC) cell line MDA-MB-231, which maintains DLC1 expression (Cao et al. 2015; Frey et al. 2022). To mimic the downregulation of DLC1 observed during tumor progression, we employed a doxycycline-inducible CRISPRoff epigenetic editing system (Nuñez et al. 2021). We specifically targeted the canonical DLC1 isoform 2, which has a well-characterized tumor suppressor function (Xue et al. 2008; Ren and Li 2021). Two independent single-guide RNAs (sgRNAs) targeting the DLC1 promoter were used, while a sgRNA targeting LacZ (sgLacZ) served as a control (Fig. 1A). Six days of doxycycline treatment reduced DLC1 transcript and protein levels by ∼50% (Fig. S1A, B; Fig. 1B, C).

**Figure 1:**
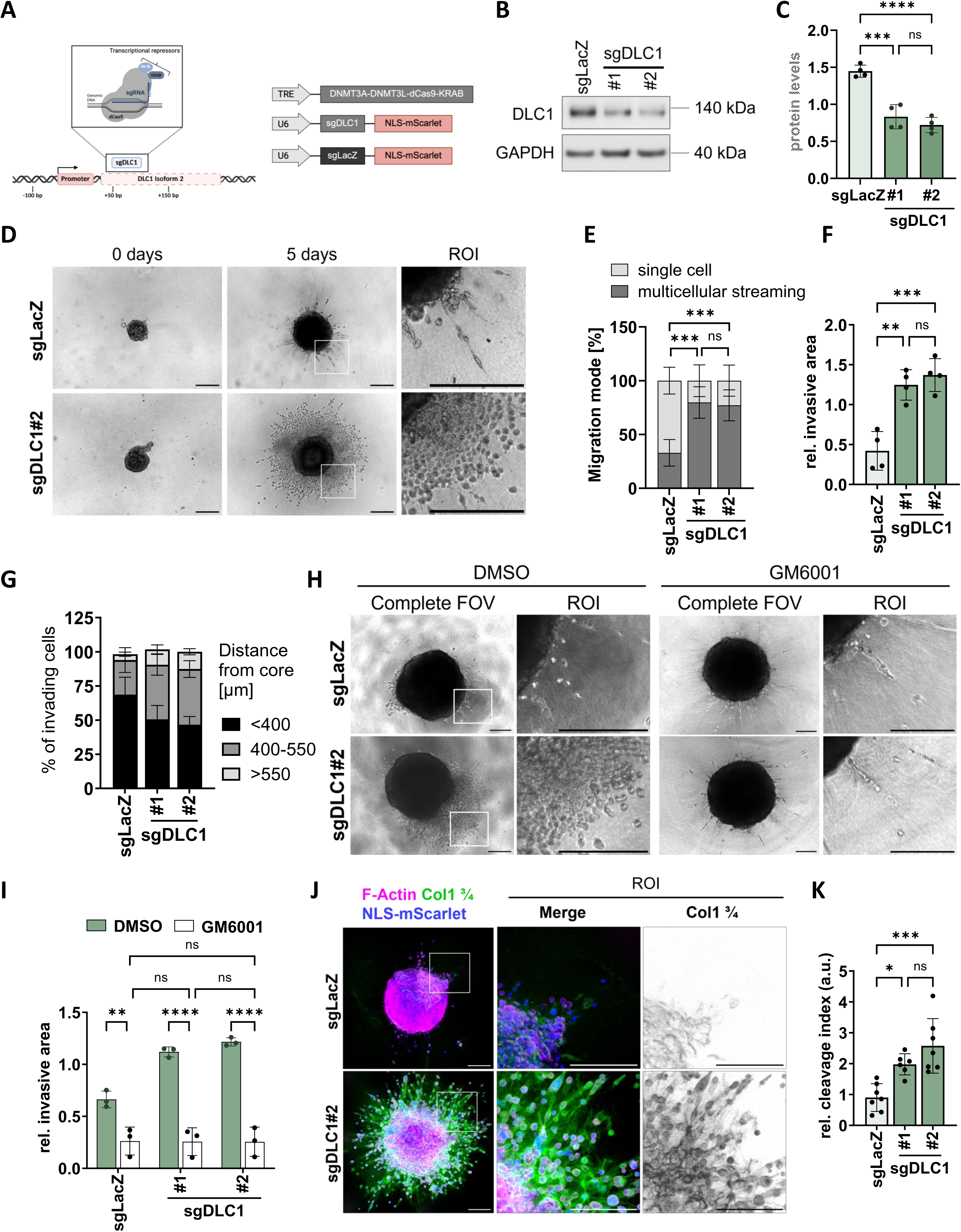
DLC1 depletion promotes spheroid invasion in 3D collagen. (A) Schematic model of the CRISPRoff strategy and sgRNAs used in this study for doxycycline inducible repression of DLC1 isoform 2. Figure was generated with Biorender.com. (B) Immunoblot analysis of whole-cell lysates from MDA-MB-231 cells expressing control (sgLacZ) or DLC1-targeting sgRNAs, following 6 days of doxycycline induction, probed for DLC1. GAPDH was used as a loading control. (C) Densitometry shows relative levels of DLC1 quantified on immunoblots representatively shown in (B) n = 4 independent experiments, data are presented as mean ± s.d.; one-way ANOVA followed by Tukey’s multiple-comparisons test (D) Representative brightfield images of doxycycline-induced MDA-MB-231 CRISPRoff spheroids expressing sgLacZ or DLC1-targeting sgRNAs (sgDLC1#2). Matched sgDLC1#1 and sgLacZ images are shown in Fig. S1C. The spheroids were embedded in 1 mg/mL collagen I, imaged on day 0 (day of collagen addition) and day 5. Boxes indicate the magnified regions of interest (ROIs). Scale bar = 400 µm (E) Quantification of invasion mode per spheroid on day 5, showing the percentage of spheroids classified as exhibiting predominantly single-cell invasion or multicellular streaming. Invasion mode was classified based on the dominant migration pattern observed per spheroid, and percentages were averaged per experiment. n = 5 independent experiments, each analyzing ≥5 spheroids per condition; data are presented as mean ± s. d.; one-way ANOVA followed by Tukey’s multiple-comparisons test. (F) Quantification of invasive area per spheroid after 5 days in collagen I based on images as representatively shown in (D). n = 4 independent experiments, each analyzing ≥5 spheroids per condition, data are presented as mean ± s.d.; one-way ANOVA followed by Tukey’s multiple-comparisons test. (G) Quantification of total invading cells and their spatial distribution relative to the spheroid core, based on representative images shown in Fig. S1F. n = 4 independent experiments, each analyzing ≥5 spheroids per condition; data are presented as mean ± s.d. Two-way repeated-measures ANOVA with sgRNA and distance category as factors, followed by Tukey’s multiple-comparisons test comparing sgRNA conditions within each distance category. For cells located <400 µm from the spheroid core, sgLacZ differed from sgDLC1#1 (P = 0.0051) and sgDLC1#2 (P = 0.0007). For cells located 400–550 µm from the core, sgLacZ differed from sgDLC1#1 (P = 0.0250) and sgDLC1#2 (P = 0.0173). No significant differences were detected between sgDLC1#1 and sgDLC1#2 or within the >550-µm category. (H) Representative brightfield images of GM6001 (50 µM) or vehicle control treated, doxycycline-induced MDA-MB-231 CRISPRoff spheroids expressing sgLacZ or DLC1-targeting sgRNAs (sgDLC1#2). Matched sgDLC1#1 and sgLacZ images are shown in Fig. S1K. The spheroids were embedded in 1 mg/mL collagen I, imaged on day 0 (day of collagen addition) and day 5. Boxes indicate the magnified regions of interest (ROIs). Scale bar = 200 µm. (I) Quantification of invasive area per spheroid after 5 days in collagen I following treatment with GM6001 (50 µM) or vehicle control, based on images representatively shown in (H). n = 3 independent experiments, each analyzing ≥5 spheroids per condition; data are presented as mean ± s.d.; two-way ANOVA with sgRNA and treatment as factors, followed by Tukey’s multiple-comparisons test. (J) Representative immunofluorescence images of doxycycline-induced MDA-MB-231 CRISPRoff spheroids expressing sgLacZ or DLC1-targeting sgRNAs (sgDLC1#2). Matched sgDLC1#1 and sgLacZ images are shown in Fig. S1L. The spheroids were embedded in 1 mg/mL collagen I, immunolabelled for MMP-cleaved collagen I (Col1-3/4; green, inverted grayscale look-up table (LUT)) and F-actin (magenta), imaged on day 5 mScarlet–NLS was used as a nuclear marker. Insets show magnified regions of interest (ROIs) of invading cell strands. Scale bar = 400 µm. (K) Quantification of collagen I cleavage (Col1-3/4 cleavage index) based on images as representatively shown in (J). n = 7 independent experiments, each analyzing ≥5 spheroids per condition; data are presented as mean ± s.d.; one-way ANOVA followed by Tukey’s multiple-comparisons test.

To determine how DLC1 regulates tumor cell invasion, we generated multicellular spheroids from the control and DLC1-depleted cells and embedded them in 1 mg/mL fibrillar collagen I gels. Time-lapse imaging of the spheroid cultures over five days revealed that DLC1 depletion triggers a striking shift in the motility phenotype (Fig. 1D, Fig. S1C). While cells from control spheroids predominantly disseminated singly into the collagen gel, cells from DLC1-depleted spheroids migrated as sheets, in which individual cells displayed highly dynamic movement while forming only short-lived cell-cell interactions (Movie 1, Fig. S1D). This phenotype resembles multicellular streaming, a migration mode described in fibrillar ECM and observed by intravital imaging in breast tumor models (Friedl and Wolf 2010; Patsialou et al. 2013; Arwert et al. 2018). Image quantification revealed that approximately 75% of DLC1-depleted spheroids exhibited a multicellular streaming phenotype compared with ∼25% of control spheroids (Fig. 1E, Fig. S1E). This was accompanied by increased invasive outgrowth, with DLC1-depleted spheroids showing a 1.9–2.2-fold increase in invasive area compared with controls (Fig. 1F, Fig. S1F). Confocal analysis also revealed a higher number of invading cells and increased invasion distance from the spheroid core (Fig. 1G, Fig. S1F, G). The observed differences were not due to changes in spheroid growth between the sgLacZ and sgDLC1 conditions during the course of the assay (Fig. S1H, I).

Cancer cells penetrate the ECM either through matrix metalloproteinase (MMP)-induced proteolytic remodeling or by squeezing through pre-existing ECM pores (Sabeh et al. 2004; Wisdom et al. 2018). To determine whether the DLC1-dependent streaming invasion program requires MMP activity, we treated the spheroid cultures with the broad-spectrum MMP inhibitor GM6001 following collagen embedding (Fig. 1H). Inhibition of MMP activity strongly reduced the invasive area of DLC1-depleted spheroids (Fig. 1I), whereas spheroid core size was unchanged (Fig. S1J, K). Consistent with MMP-dependent collagen remodeling, immunolabeling for MMP-mediated collagen I cleavage sites (Col1 3/4) (Haeger et al. 2014) revealed a strong and significant increase in collagen degradation surrounding DLC1-depleted spheroids when compared with controls (Fig. 1J, K; Fig. S1L). Together, these results indicate that DLC1 depletion promotes invasion into 3D collagen matrices via an MMP-dependent multicellular streaming mode.

### DLC1 depletion stabilizes polarized RhoA activity to drive actomyosin-dependent multicellular streaming

RhoA-driven actomyosin contractility supports force generation and invasive movement through 3D matrices (Friedl and Alexander 2011; Li et al. 2011; Wang et al. 2020). However, the contribution of the Rho pathway specifically to multicellular streaming remains incompletely understood. We therefore tested whether DLC1 depletion promotes streaming invasion through enhanced actomyosin contractility. Consistent with its role as a negative regulator of Rho signaling (Wong et al. 2005), DLC1-depleted cells showed elevated F-actin signals in 3D (Fig. S2A, B). While inhibition of myosin II activity with blebbistatin did not affect the motility of the sgLacZ controls, it reduced the invasion of DLC1-depleted spheroids (Fig. 2A, B; Fig. S2C), fully abolishing their multicellular streaming phenotype (Fig. 2C). This occurred despite persistent collagen cleavage (Fig. S2D). Blebbistatin treatment did not affect the spheroid core size (Fig. S2E). Together, these data indicate that actomyosin contractility is required for the multicellular streaming phenotype of the DLC1-depleted cells.

**Figure 2:**
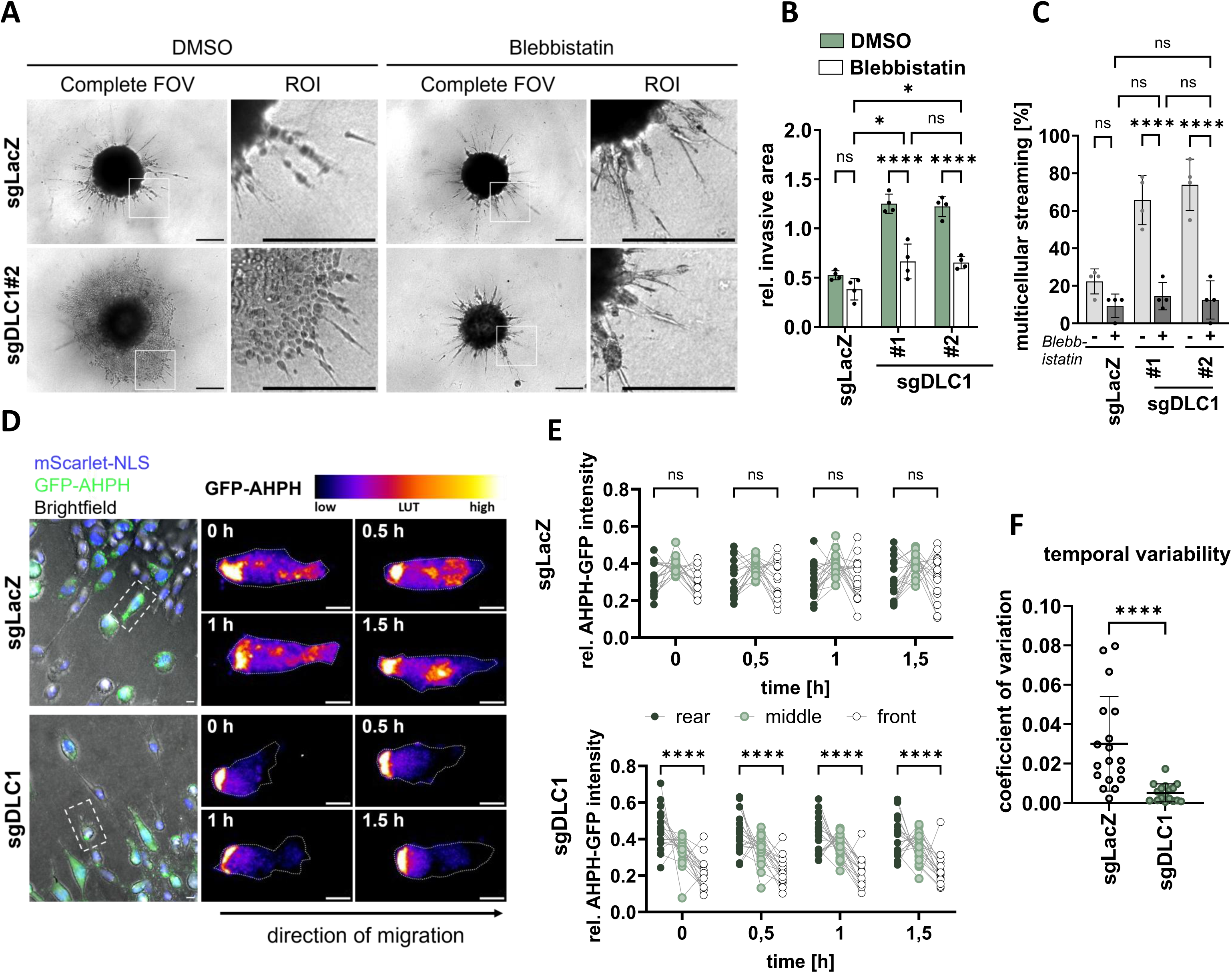
DLC1 depletion promotes contractility-dependent multicellular streaming and stabilizes Rho-GTP polarity. (A) Representative brightfield images of blebbistatin (10 µM) or vehicle control treated, doxycycline-induced MDA-MB-231 CRISPRoff spheroids expressing sgLacZ or DLC1-targeting sgRNAs (sgDLC1#2). Matched sgDLC1#1 and sgLacZ images are shown in Fig. S2C. The spheroids were embedded in 1 mg/mL collagen I and were imaged at day 0 (day of collagen addition) and day 5. Boxes indicate the magnified regions of interest (ROIs). Scale bar = 400 µm. (B) Quantification of invasive area per spheroid after 5 days in collagen I following treatment with blebbistatin (10 µM) or vehicle control, based on images as representatively shown in (A). n = 4 independent experiments, each analyzing ≥5 spheroids per condition; data are presented as mean ± s.d.; two-way ANOVA with sgRNA and treatment as factors, followed by Tukey’s multiple-comparisons test. (C) Quantification of invasion mode per spheroid at day 5 following treatment with blebbistatin (10 µM) or vehicle control, based on images as representatively shown in (A). Invasion mode was classified based on the dominant migration pattern observed per spheroid, and percentages were averaged per experiment. n = 4 independent experiments, each analyzing ≥5 spheroids per condition; data are presented as mean ± s.d.; two-way ANOVA with sgRNA and treatment as factors, followed by Tukey’s multiple-comparisons test. (D) Representative maximum-intensity projections from time-lapse live-cell imaging of doxycycline-induced MDA-MB-231 CRISPRoff spheroids expressing sgLacZ or DLC1-targeting sgRNAs together with the independent expression of the Rho activity biosensor GFP–AHPH-WT (green). The spheroids were embedded in 1 mg/mL collagen I and time-lapse imaging was performed on day 5 after collagen addition at 0, 0.5, 1 and 1.5 h. Nuclei are shown in blue. GFP–AHPH-WT signal is additionally displayed using a pseudocolor LUT, with pixel intensities ranging from 0 (minimum) to 255 (maximum). Scale bar = 10 µm. (E) Distribution of GFP-AHPH WT biosensor signal in invading cells over time. Relative GFP–AHPH intensity was quantified in rear, middle, and front regions of invading cells at the indicated time points and expressed as a fraction of the total cellular signal. Data are presented as mean ± s.d. from 18 cells pooled across three independent experiments. Each dot represents one cell. sgDLC1#1 and sgDLC1#2 were pooled for analysis. Statistical testing by one-way ANOVA followed by Tukey’s multiple-comparisons test. (F) Temporal variability of RhoA-GTP polarity quantified as the coefficient of variation (CV) of the GFP–AHPH front/rear polarity ratio over time. CV values were calculated per cell from front/rear ratios measured at 0, 0.5, 1 and 1.5 h. Data are presented as mean ± s.d. from 18 cells pooled across three independent experiments. Each dot represents one cell; sgDLC1#1 and sgDLC1#2 were pooled for analysis. Error bars indicate mean ± s.d., statistical significance was determined using a two-tailed Mann–Whitney U-test.

We next investigated the subcellular distribution of active Rho in cells invading in the 3D environment. To this end, we employed a stable cell line expressing the GFP–AHPH biosensor, an established localization-based reporter of endogenous Rho activity dynamics (Piekny and Glotzer 2008; Priya et al. 2015; Gaston et al. 2021). Immunoblotting of cell lysates confirmed full-length GFP–AHPH expression without detectable degradation products (Fig. S2F). Knockdown of individual Rho family members furthermore showed that in our model system the sensor predominantly reported on RhoA and RhoB activity (Fig. S2G, H). Importantly, biosensor expression did not affect the cell motility phenotype in spheroid experiments (Fig. S2I, J).

Live imaging of cells invading from the spheroid core into the surrounding collagen matrix revealed rearward enrichment of the GFP–AHPH sensor (Fig. 2D), consistent with the association of RhoA in actomyosin-driven tail retraction (Petrie et al. 2009; Jia et al. 2026). Interestingly, while control cells exhibited Rho-GTP pools, which dynamically redistributed during migration, DLC1-depleted cells displayed pronounced and persistent front–rear polarity with stabilized rearward Rho-GTP enrichment (Fig. 2D). Quantitative analysis confirmed increased spatial asymmetry of Rho-GTP localization (Fig. 2E) together with reduced temporal variability of signal distribution in the DLC1-depleted cells (Fig. 2F).

Together, these data indicate that DLC1 depletion promotes efficient actomyosin-dependent multicellular streaming by stabilizing Rho activity gradients.

### DLC1 depletion promotes invasion in collagen ECM by increasing β1 integrin abundance

While imaging the spheroids, we observed that collagen fibers reorganized and formed tracks around invading cells, being most pronounced in areas of multicellular streams (Fig. S3A). This suggests that increased matrix engagement contributes to the invasive behavior of the DLC1 depleted cells. Because traction-dependent invasion within fibrillar ECM relies on β1 integrins as the main collagen receptors (Sabeh et al. 2004; Friedl and Wolf 2010; Chastney et al. 2025), we investigated the contribution of β1 integrin to the invasive behavior. Blocking β1 integrin with the AIIB2 antibody (Berrazouane et al. 2019; Park et al. 2008) induced receptor internalization without affecting spheroid growth (Fig. S3B, C, E). Notably, this treatment fully abolished the coordinated streaming seen in DLC1-depleted spheroids. Instead, cells formed rounded, compact clusters with strongly limited displacement into the surrounding matrix (Fig. 3A, B; Fig. S3D). This morphology, which was also observed in sgLacZ cells, is consistent with impaired force transmission to the ECM and reduced traction-dependent migration (Wolf et al., 2007; Friedl and Alexander, 2011).

**Figure 3:**
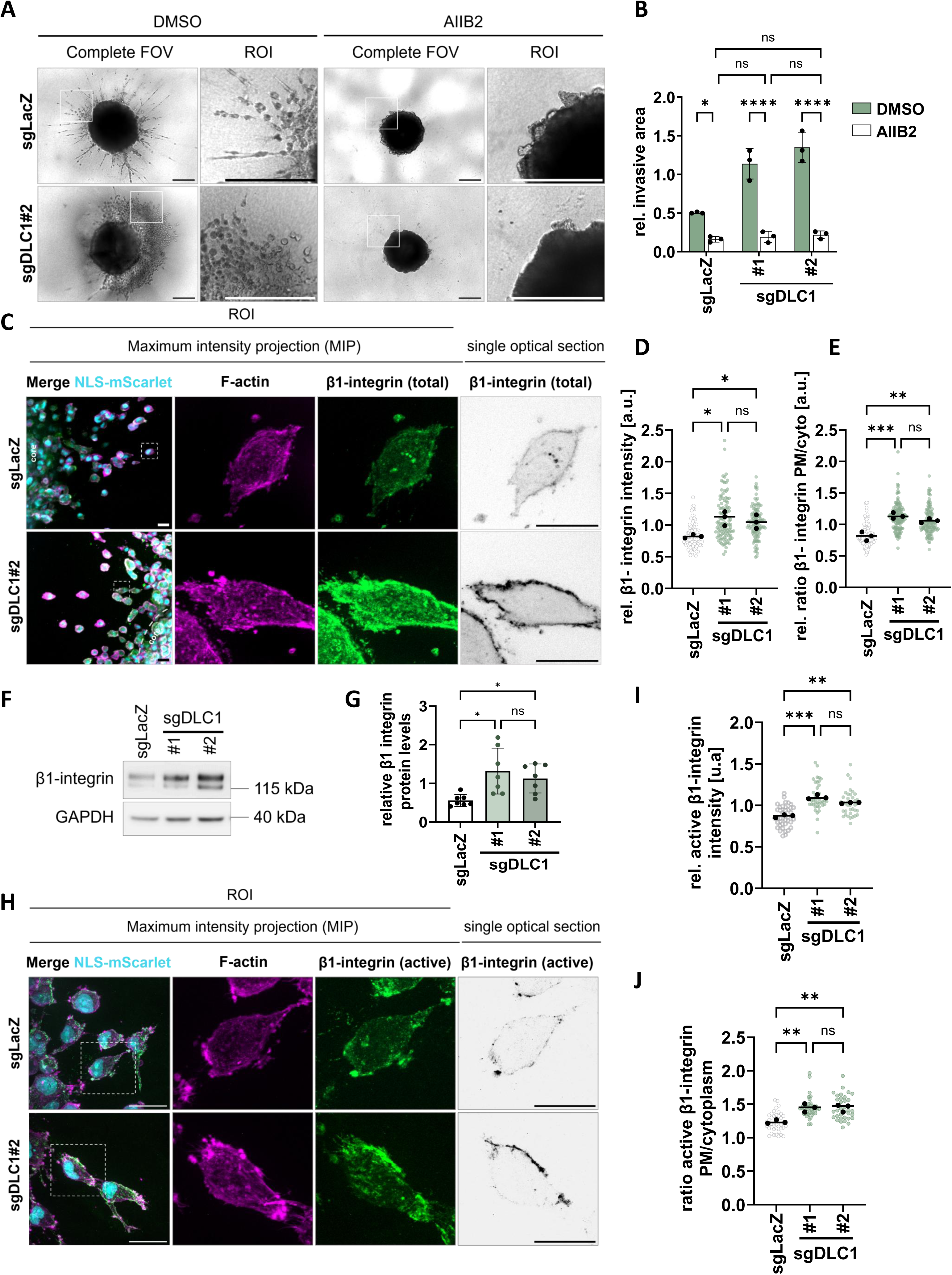
β1 integrin function is required for multicellular streaming and is spatially reorganized upon DLC1 depletion. (A) Representative bright-field images of AIIB2 (1 µg/ml) or vehicle control treated doxycycline-induced MDA-MB-231 CRISPRoff spheroids expressing sgLacZ or DLC1-targeting sgRNAs (sgDLC1#2). Matched sgDLC1#1 and sgLacZ images are shown in Fig. S3D. The spheroids were embedded in 1 mg/mL collagen I, imaged on day 5. Boxes indicate the magnified regions of interest (ROIs). Scale bar = 400 µm. (B) Quantification of invasive area per spheroid after 5 days in collagen I following treatment with AIIB2 (1 µg/ml) or vehicle control based on images as representatively shown in (A). n = 3 independent experiments, each analyzing ≥5 spheroids per condition. Data are presented as mean ± s.d.; two-way ANOVA with sgRNA and treatment as factors, followed by Tukey’s multiple-comparisons test. (C) Representative immunofluorescence images of invading cells from doxycycline-induced MDA-MB-231 CRISPRoff spheroids expressing sgLacZ or DLC1-targeting sgRNAs (sgDLC1#2). The spheroids were embedded in 1 mg/mL collagen I, stained for total β1 integrin (green, grayscale LUT) and F-actin (magenta) on day 5. Shown are single optical sections for β1 integrin and maximum-intensity projections (MIPs) are shown for merged images. Invading cells from sgLacZ and DLC1-depleted spheroids are shown. Scale bar = 30 µm. (D) Quantification of total cellular β1 integrin fluorescence intensity per invading cell based on images as representatively shown in (C). n = 3 independent experiments, each analyzing ≥ 20 cells per condition. Data are presented as mean ± s.d.; one-way ANOVA followed by Tukey’s multiple-comparisons test. (E) Quantification of the plasma membrane-to-cytoplasmic β1 integrin fluorescence intensity ratio in invading cells based on images as representatively shown in (C). n = 3 independent experiments, each analyzing ≥ 40 cells per condition. Data are presented as mean ± s.d.; one-way ANOVA followed by Tukey’s multiple-comparisons test. (F) Immunoblot analysis from whole-cell lysates of MDA-MB-231 cells expressing control (sgLacZ) or DLC1-targeting sgRNAs, following 6 days of doxycycline induction and cultured on collagen-coated dishes. Probed for β1 integrin. GAPDH was used as loading control. (G) Densitometry shows relative levels from β1 integrin quantified on immunoblots representatively shown in (F). n = 7 independent experiments, data are presented as mean ± s.d.; one-way ANOVA followed by Tukey’s multiple-comparisons test. (H) Representative immunofluorescence images of invading cells from doxycycline-induced MDA-MB-231 CRISPRoff spheroids expressing sgLacZ or DLC1-targeting sgRNAs (sgDLC1#2). The spheroids were embedded in 1 mg/mL collagen I, stained for active β1 integrin (green, grayscale LUT) and F-actin (magenta) on day 5. Shown are single optical sections for active β1 integrin and maximum intensity projections (MIPs) for merged images. Invading cells from sgLacZ and DLC1-depleted spheroids are shown. Scale bar = 30 µm (I) Quantification of total cellular active β1 integrin fluorescence intensity per invading cell based on images as representatively shown in (H). n = 3 independent experiments, each analyzing ≥ 15 cells per condition. Data are presented as mean ± s.d.; one-way ANOVA followed by Tukey’s multiple-comparisons test. (J) Quantification of plasma membrane enrichment of active β1 integrin in invading cells based on images as representatively shown in (H), expressed as plasma membrane-to-cytoplasmic fluorescence intensity ratio. n = 3 independent experiments, each analyzing ≥ 15 cells per condition. Data are presented as mean ± s.d.; one-way ANOVA followed by Tukey’s multiple-comparisons test.

To explore in more detail how β1 integrin might be regulated by DLC1, we next examined β1 integrin organization by immunofluorescence analysis. Total β1 integrin levels and the plasma membrane-to-cytoplasmic ratio were both significantly increased in DLC1-depleted cells (Fig. 3C-E). These results were confirmed by immunoblotting, which showed increased total β1 integrin protein abundance in sgDLC1 cells (Fig. 3F, G). Furthermore, active β1 integrin, detected using the conformation-sensitive antibody 12G10 (Humphries et al. 2005), exhibited increased plasma membrane enrichment in DLC1-depleted cells and displayed clear spatial polarization at sites of matrix engagement within invading cells (Fig. 3H-J). No changes in β1 integrin expression were detected by RT-qPCR (Fig. S3F), indicating that DLC1 regulates β1 integrin abundance and localization post-transcriptionally. Collectively, these results demonstrate that DLC1 downregulation increases β1 integrin abundance and activation, thereby promoting invasion through collagen-rich ECM.

### DLC1 re-expression suppresses multicellular streaming in a GAP-dependent manner

We next asked whether the invasion mode induced by engineered DLC1 depletion is reflected in a metastatic breast cancer model that has intrinsically lost DLC1 expression. To this end, we analyzed the bone-tropic MDA-MB-231 subline BoM-1833 (Kang et al. 2003; Cox et al. 2015), a bone-metastatic derivative of MDA-MB-231 cells that showed markedly reduced DLC1 expression (Fig. 4A, Fig. S4A). In contrast to the mainly single-cell invasion mode of the parental MDA-MB-231 spheroids, BoM-1833 spheroids showed a pronounced multicellular streaming phenotype reminiscent of the DLC1-depleted cells (∼80% vs. ∼20% of controls; Fig. 4B, C), together with a ∼2.5–3-fold increase in invasive area (Fig. 4D). Indeed, streaming of the BoM-1833 spheroids required MMP activity, actomyosin contractility and β1 integrin function, as treatment with GM6001, blebbistatin and AIIB2, respectively, abolished the invasive streaming program and eliminated differences between MDA-MB-231 and BoM-1833 spheroids (Fig. S4B, C).

**Figure 4:**
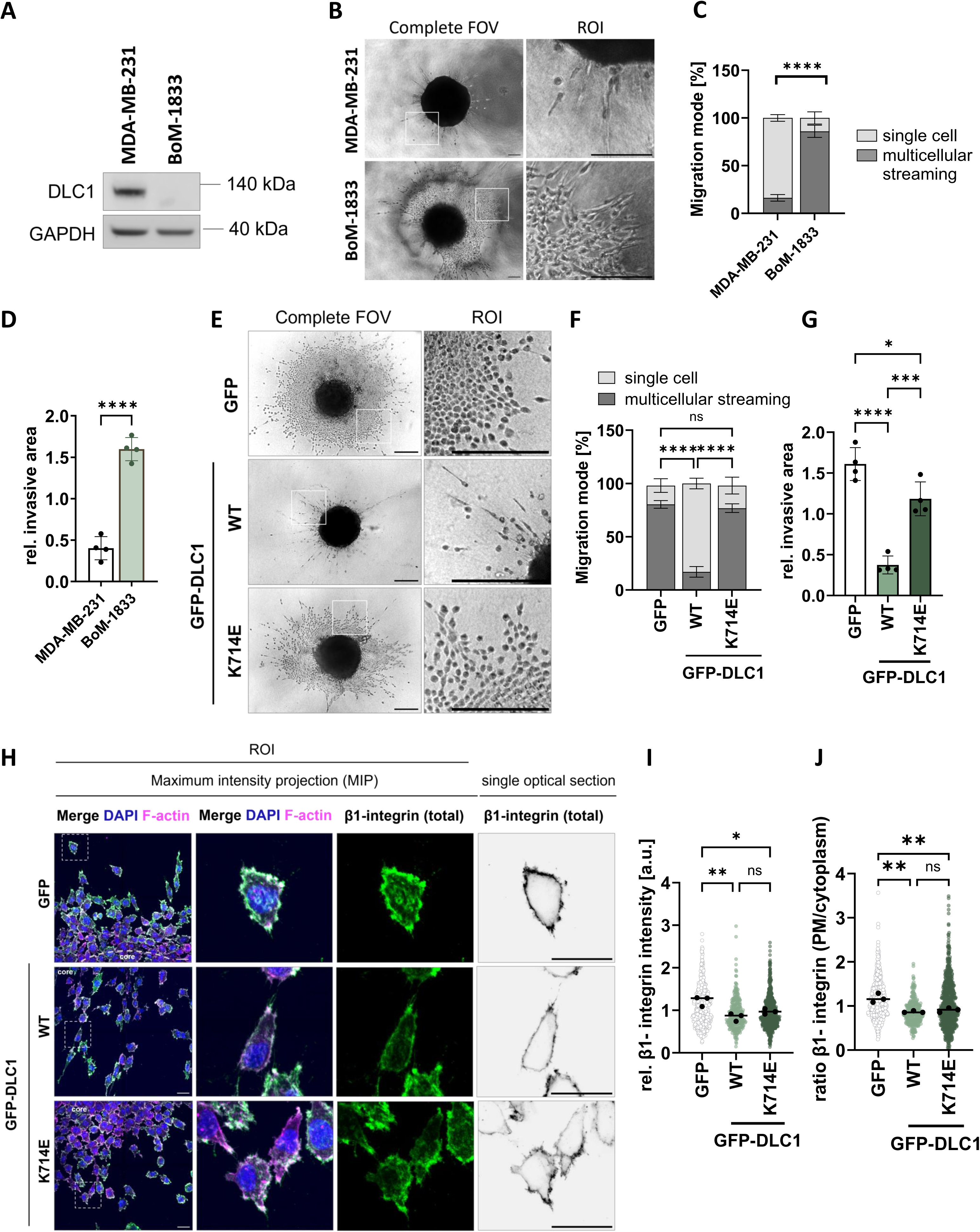
DLC1 re-expression constrains multicellular streaming and β1 integrin enrichment in a bone-metastatic MDA-MB-231 subline. (A) Immunoblot analysis of whole-cell lysates of parental MDA-MB-231 and bone-tropic BoM-1833 cells, probed for DLC1. GAPDH was used as loading control. (B) Representative brightfield images from MDA-MB-231 and BoM-1833 spheroids. The spheroids were embedded in 1 mg/ml collagen, imaged after 5 days of invasion. Boxes indicate the magnified regions of interest (ROIs). Scale bar = 400 µm. (C) Quantification of invasion mode per spheroid at day 5 showing the percentage of MDA-MB-231 and BoM-1833 spheroids exhibiting multicellular streaming versus predominantly single-cell invasion, based on images as representatively shown in (B). Invasion mode was classified based on the dominant migration pattern observed per spheroid, and percentages were averaged per experiment. n = 4 independent experiments, each analyzing ≥5 spheroids per condition; data are presented as mean ± s.d.; one-way ANOVA followed by Tukey’s multiple-comparisons test. (D) Quantification of invasive outgrowth area per spheroid after 5 days in collagen I based on images as representatively shown in (B). n = 4 independent experiments, each analyzing ≥5 spheroids per condition. Data are presented as mean ± s.d.; unpaired two-tailed t-test with Welch’s correction. (E) Representative brightfield images of doxycycline-induced BoM-1833 spheroids expressing GFP control, GFP–DLC1 (WT), or GFP–DLC1(K714E). The spheroids were embedded in 1 mg/mL collagen I, imaged at day 0 (day of collagen addition) and day 5. Boxes indicate annotated the magnified regions of interest (ROIs). Scale bar = 400 µm. (F) Quantification of invasion mode per spheroid on day 5, showing the percentage of spheroids classified as exhibiting predominantly single-cell invasion or multicellular streaming, based on images as representatively shown in (E). Invasion mode was classified based on the dominant migration pattern observed per spheroid, and percentages were averaged per experiment. Data presented as mean ± s.d. from n = 5 independent experiments, each analyzing ≥5 spheroids per condition. One-way ANOVA followed by Tukey’s multiple-comparisons test. (G) Quantification of invasive area per spheroid after 5 days in collagen I based in images as representatively shown in (E). n = 4 independent experiments, analyzing ≥5 spheroids per condition. Data are presented as mean ± s.d.; one-way ANOVA followed by Tukey’s multiple-comparisons test. (H) Representative immunofluorescence images of invading cells from doxycycline-induced BoM-1833 spheroids expressing GFP control, GFP–DLC1 (WT), or GFP–DLC1(K714E). The spheroids were embedded in 1 mg/mL collagen I, stained for total β1 integrin (green, grayscale LUT) and F-actin (magenta) and imaged on day 5 after invasion. Shown are single optical sections for β1 integrin and maximum-intensity projections (MIPs) for merged images. Invading cell from BoM-1833 spheroids expressing GFP, GFP–DLC1, or GFP–DLC1(K714E) are shown. Scale bar = 30 µm. (I) Quantification of total cellular β1 integrin fluorescence intensity per invading cell based on images as representatively shown in (H). n = 3 independent experiments, each analyzing ≥ 20 cells per condition. Data are presented as mean ± s.d.; one-way ANOVA followed by Tukey’s multiple-comparisons test. (J) Quantification of the plasma membrane-to-cytoplasmic β1 integrin fluorescence intensity ratio in invading cells as representatively shown in (H). n = 3 independent experiments, each analyzing ≥ 20 cells per condition. Data are presented as mean ± s.d.; one-way ANOVA followed by Tukey’s multiple-comparisons test.

To test whether DLC1 and its RhoGAP activity is sufficient to suppress the BoM-1833 streaming phenotype, we generated cell lines with stable, doxycycline-inducible expression of wild-type or GAP-inactive (K714E) DLC1 (Qian et al. 2007; Wong et al. 2005). A cell line expressing unconjugated GFP was used as a control. Comparable expression of the DLC1 variants was confirmed by immunoblotting (Fig. S4D). Localization to focal adhesions was confirmed by imaging of two-dimensional cultures (Fig. S4E). Notably, re-expression of wild-type DLC1 reduced the percentage of spheroids with a streaming phenotype from ∼80% in GFP controls to ∼20% (Fig. 4E, F). The invasive outgrowth area was decreased by approximately 3-fold (Fig. 4G). In contrast, the GAP-deficient mutant DLC1(K714E) failed to suppress the streaming behavior (Fig. 4F), although it induced a modest (∼1.2 fold) but significant reduction in the invasive area (Fig. 4G).

We next examined the levels and organization of β1 integrin. Mirroring the findings in the sgDLC1 system, immunofluorescence analyses showed that expression of wild-type DLC1 reduced the abundance of the total β1 integrin protein relative to GFP controls (Fig. 4H-J,). Intriguingly, in immunofluorescence stainings of DLC1(K714E)-expressing cells, β1 integrin protein levels and distribution were largely comparable to those in wild-type DLC1–expressing cells (Fig. 4H-J). Similarly, immunoblotting of cell lysates revealed decreased total β1 integrin protein abundance in both wild-type DLC1 and DLC1(K714E) expressing cells (Fig. S4F, G). Thus, whereas DLC1 suppresses the streaming invasion mode through its RhoGAP activity, additional GAP-independent functions appear to contribute to the regulation of β1 integrin function and overall invasion.

### DLC1 limits β1 integrin–dependent invasion through coordinated activity of its RhoGAP and LD-like domains

To identify a potential GAP-independent function, we focused on the DLC1 LD-like motif. This motif was shown to mediate the interaction of DLC1 with the R8 domain of talin (Li et al. 2011; Zacharchenko et al. 2016). Talin binding to integrin β cytoplasmic tails drives integrin inside-out activation and clustering at sites of matrix engagement (Calderwood 2004; Lee et al. 2009; Moreno-Layseca et al. 2019). We therefore hypothesized that talin binding might be involved in the regulation of β1 integrin by DLC1.

To test this, we generated BoM-1833 cells with inducible expression of a GAP-inactive GFP-DLC1 mutant lacking the LD motif (K714E/ΔLD) and confirmed loss of talin binding by co-immunoprecipitation (Fig. 5A). As expected, in spheroid assays, the migration mode of the GFP-DLC1(K714E/ΔLD) cells was comparable to that of the GFP controls (Fig. 5B, C). Strikingly, combined disruption of GAP activity and talin binding fully abolished the ability of DLC1 to suppress invasion (Fig. 4G vs. Fig. 5D). Importantly, GFP-DLC1(K714E/ΔLD) failed to reduce β1 integrin abundance and plasma membrane localization, which remained at levels comparable to those observed in GFP control cells (Fig. 5E–G, Fig. S5A, B). Together, these results support a model in which DLC1 restrains β1 integrin localization and collagen invasion through coordinated RhoGAP activity and LD-dependent talin binding.

**Figure 5:**
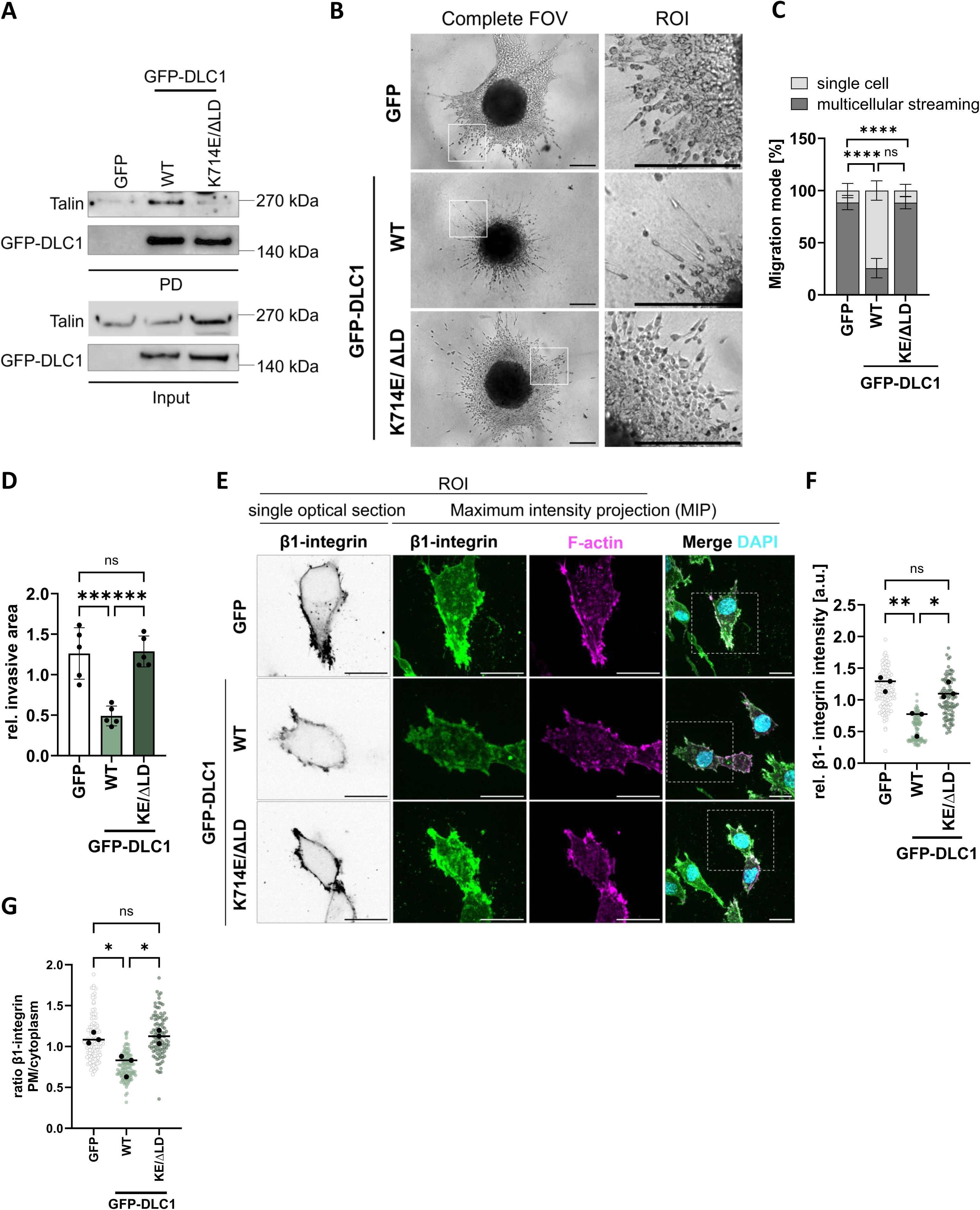
DLC1 limits β1 integrin–dependent invasion through coordinated RhoGAP activity and its LD-binding motif. (A) Immunoblot analysis of whole-cell lysates and GFP pulldowns from BoM-1833 cells expressing GFP control, GFP–DLC1 (WT), or GFP–DLC1 (K714E/ΔLD), following 3 days of doxycycline induction, probed for GFP and talin. (B) Representative brightfield images of doxycycline-induced BoM-1833 spheroids expressing GFP control, GFP–DLC1 (WT), or GFP–DLC1 (K714E/ΔLD). The spheroids were embedded in 1 mg/mL collagen I and imaged after 5 days. Boxes indicate the magnified regions of interest (ROIs). Scale bar = 400 µm. (C) Quantification of invasion mode per spheroid on day 5, showing the percentage of spheroids classified as exhibiting predominantly single-cell invasion or multicellular streaming based on images as representatively shown in (B). Invasion mode was classified based on the dominant migration pattern observed per spheroid, and percentages were averaged per experiment. Data presented as mean ± s.d. from n = 5 independent experiments, each analyzing ≥5 spheroids per condition. One-way ANOVA followed by Tukey’s multiple-comparisons test. (D) Quantification of invasive area per spheroid after 5 days in collagen I based on images as representatively shown in (B) Data presented as mean ± s.d. from n = 5 independent experiments, each analyzing ≥5 spheroids per condition. One-way ANOVA followed by Tukey’s multiple-comparisons test. (E) Representative cropped regions showing cells within invasive sheets emerging from doxycycline-induced BoM-1833 spheroid cores expressing GFP, GFP–DLC1, or GFP–DLC1(K714E/ΔLD). Spheroids were embedded in 1 mg/mL collagen I, allowed to invade for 5 days, and stained for total β1 integrin (green; shown separately using a grayscale LUT) and F-actin (magenta). Single optical sections are shown for β1 integrin, whereas merged images are displayed as maximum-intensity projections. Scale bar, 20 µm. (F) Quantification of total cellular β1 integrin fluorescence intensity per invading cell based on images as representatively shown in (E). n = 3 independent experiments, each analyzing ≥ 15 cells per condition. Data are presented as mean ± s.d.; one-way ANOVA followed by Tukey’s multiple-comparisons test. (G) Quantification of plasma membrane enrichment of β1 integrin in invading cells as representatively shown in (E), expressed as plasma membrane-to-cytoplasmic fluorescence intensity ratio. n = 3 independent experiments, each analyzing ≥ 15 cells per condition. Data are presented as mean ± s.d.; one-way ANOVA followed by Tukey’s multiple-comparisons test.

### The DLC1-controlled β1 integrin–dependent invasion program is conserved in patient-derived metastatic organoids

To determine whether the DLC1-controlled invasion program identified in the MDA-MB-231 cell lines is conserved in primary cells, we used a patient-derived breast cancer organoid model established from pleural effusion (Önder et al. 2023), that retained DLC1 expression (Fig. 6A). DLC1 was downregulated using the doxycycline-inducible CRISPRoff epigenetic editing system and efficient repression of the DLC1 isoform 2 was confirmed by qPCR after six days of doxycycline treatment (Fig. 6A). Following transduction, the organoids retained a normal and uniform morphology (Fig. S6A).

**Figure 6:**
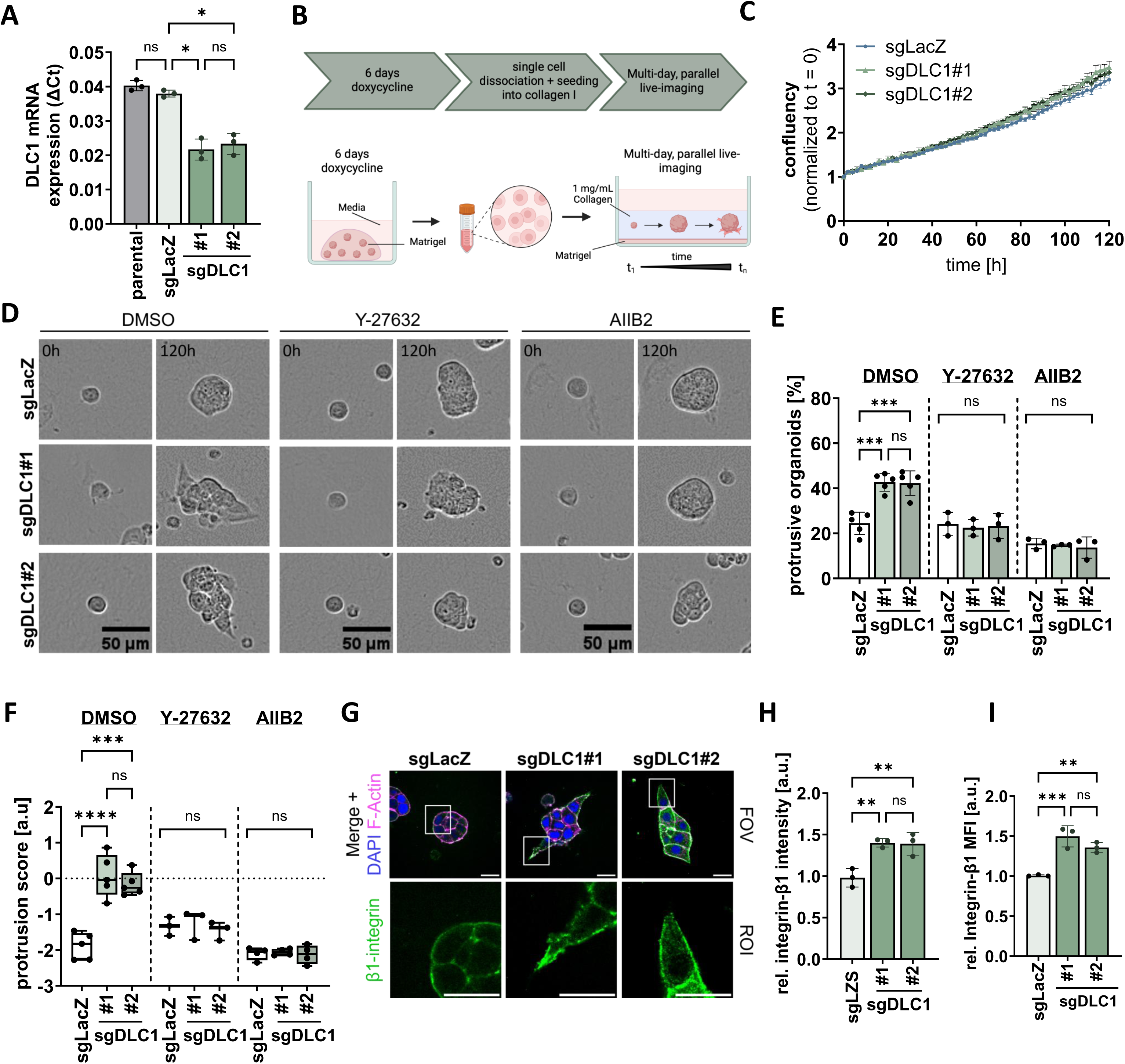
DLC1 regulates a conserved β1 integrin–dependent invasion program in patient-derived metastatic organoids. (A) Quantitative PCR analysis of DLC1 isoform 2 mRNA levels in breast cancer CRISPRoff organoids expressing control (sgLacZ) or DLC1-targeting sgRNAs, following 6 days of doxycycline induction. Parental breast cancer organoids were included as control. n = 3 independent experiments, data are presented as mean ± s.d.; one-way ANOVA one-way ANOVA followed by Tukey’s multiple-comparisons test. (B) Experimental workflow for organoid invasion assays. Patient-derived breast cancer organoids were cultured in Matrigel and induced with doxycycline for 6 days to establish CRISPRoff-mediated DLC1 repression. Organoids were then dissociated into single cells, embedded in collagen I matrices and monitored by live-cell imaging for 5 days prior to endpoint analysis. Figure was generated with Biorender.com. (C) Relative NLS-mScarlet fluorescence intensity in doxycycline-induced CRISPRoff organoids expressing sgLacZ or DLC1-targeting sgRNAs. Fluorescence intensity was used as a measure of organoid growth and normalized to t = 0. The organoids were embedded in 1 mg/mL collagen I and imaged for 5 days. Data represent mean ± s.d. from n = 1 independent experiments, each analyzing ≥50 organoids per condition in one experiment. (D) Representative magnified brightfield images of Y-27632 (50 µM), AIIB2 (1 µg/ml) and vehicle control treated doxycycline-induced CRISPRoff organoids expressing sgLacZ or DLC1-targeting sgRNAs. The organoids were embedded in 1 mg/mL collagen and imaged after 5 days. Scale bar = 50 µm. (E, F) Quantification of organoid morphology showing (E) the percentage of organoids classified as protrusive and (F) the protrusion score based on shape classifiers. Data are presented as mean ± s.d. from n = 5 independent experiments, each analyzing ≥50 organoids per condition and experiment. one-way ANOVA followed by Tukey’s multiple-comparisons test. (G) Representative immunofluorescence images of doxycycline-induced CRISPRoff organoids expressing sgLacZ or DLC1-targeting sgRNAs, stained for total β1 integrin (green) and F-actin (magenta). Nuclei were stained with DAPI (blue). Shown are single optical sections. Boxes indicate the magnified regions of interest (ROIs). Scale bar = 20 µm. (H) Quantification of total cellular β1 integrin fluorescence intensity per organoid in conditions based on images as representatively shown in (G). Data are presented as mean ± s.d. from n = 5 independent experiments, each analyzing ≥10 organoids per condition and experiment. One-way ANOVA followed by Tukey’s multiple-comparisons test. (I) Flow cytometric analysis of cell-surface β1 integrin levels in non-permeabilized cells isolated from collagen-embedded organoids expressing control (sgLacZ) or DLC1-targeting sgRNAs. Data are presented as mean ± s.d. from n = 3 independent experiments; one-way ANOVA followed by Tukey’s multiple-comparisons test.

We next dissociated the organoids grown in matrigel domes, embedded the cells in 1 mg/mL fibrillar collagen and monitored the growth and morphology of the 3D cultures over five days in the presence of 2 µg/ml doxycycline ((Nikolatou et al. 2023), Fig. 6B). DLC1 depletion did not affect organoid growth under these conditions (Fig. 6C, S6C). Notably, while control organoids predominantly maintained a rounded morphology with little protrusive activity, DLC1-repressed organoids frequently extended invasive protrusions into the surrounding matrix (Fig. 6D, Movie 2).

Quantification of this phenotype revealed a marked increase in the proportion of protrusive organoids to ∼40–50% for the sgDLC1 condition compared with ∼20–25% in controls. To quantify organoid protrusiveness, we calculated a composite protrusion score based on circularity, aspect ratio and solidity. DLC1 downregulation increased this score by approximately two-fold (Fig. 6E, F). To test whether this phenotype depends on the mechanisms identified in the cell lines, we inhibited actomyosin contractility and β1 integrin function, respectively. ROCK inhibition by Y-27632 enforced a rounded morphology in both control and DLC1-repressed organoids and abolished the differences in protrusion frequency and score. Similarly, β1 integrin blockade using AIIB2 suppressed protrusion formation and eliminated the DLC1-dependent increase in invasiveness (Fig. 6D–F).

We next examined β1 integrin regulation in the organoids cultured in collagen matrices. In agreement with the cell line data, immunostaining revealed increased β1 integrin abundance in the DLC1-depleted organoids, with prominent enrichment along invasive protrusions (Fig. 6G, H). Consistently, immunoblotting showed increased total β1 integrin levels (Fig. S6D, E), and a flow cytometry protocol established for fixed cells isolated from the collagen matrices confirmed elevated surface β1 integrin levels upon DLC1 depletion (Fig. 6I). Together, these data demonstrate that DLC1 repression promotes a protrusive, β1 integrin– and contractility-dependent invasion program in patient-derived metastatic organoids, establishing DLC1 as a conserved regulator of β1 integrin–dependent 3D invasion.

## Discussion

Cell invasion through 3D ECM is a highly coordinated process in which cells sense, respond to, and remodel their surrounding matrix (Friedl and Wolf 2010; Paul et al. 2017). Although this process is central to normal development and deregulated in cancer, the molecular mechanisms that coordinate it remain incompletely understood. Here, we identify DLC1, a tumor suppressor frequently deleted or silenced in cancer (Xue et al. 2008; Wang et al. 2014), as a key regulator of invasive behavior in collagen-rich matrices, acting through two cooperative, mechanistically distinct arms: GAP-dependent control of Rho signaling and contractility, and LD-like motif-dependent regulation of β1-integrin function.

Using epigenetically edited MDA-MB-231 spheroid cultures embedded in collagen, we show that DLC1 downregulation promotes a multicellular streaming phenotype, increasing cancer cell dissemination into the surrounding ECM. Conversely, DLC1 re-expression in BoM-1833 cells, a bone-metastatic MDA-MB-231 subline with low endogenous DLC1 (Wang et al. 2014), fully suppressed this invasion program. Together, these complementary models converge on DLC1 as a molecular brake on streaming-based cell invasion. This motility mode has been previously observed by intravital imaging in breast carcinomas and, while mechanistically poorly understood, is thought to provide cancer cells with a more efficient dissemination route by allowing cells to share paths, coordinate matrix remodeling and amplify force generation (Arwert et al. 2018; Patsialou et al. 2013). We furthermore found that downregulation of DLC1 in patient-derived breast cancer organoids promoted aberrant protrusive behavior in a 3D collagen environment, which was sensitive to inhibition of β1 integrin function and Rho-ROCK signaling. These findings, provide evidence for a conserved multifaceted role of DLC1 in primary models beyond established cell lines.

The dependence of multicellular streaming on the RhoGAP activity of DLC1 suggests that altered Rho regulation is a central driver of this phenotype. Consistent with this, DLC1 depletion promoted prominent stabilization of rear-polarized Rho-GTP during invasion, most likely corresponding to active RhoA, as detected with the AHPH Rho activity biosensor. These data suggest that DLC1 downregulation not only increases global RhoA activity, but also alters where active RhoA is positioned and maintained during invasion. This spatial organization may explain why DLC1-low cells form persistent multicellular streams, as rear actomyosin contractility can drive 3D invasion by generating traction against the matrix (Poincloux et al. 2011). These traction forces are known to be transmitted through cell adhesions, where local GTPase control amplifies force generation by feedback mechanisms (Kanchanawong and Calderwood 2023). Consistently, Müller et al. showed that adhesion sites position distinct GEFs and GAPs to generate local GTPase activity zones, supporting the broader view that adhesion proteins control Rho-family signaling in space and time and in relation to ECM engagement (Müller et al. 2020).

Further studies underscore this spatial model of Rho regulation. In neural crest migration, DLC1 shapes front–rear RhoA polarity with consequences for directional movement (Liu et al. 2017), and during neurite outgrowth, DLC1 and p190RhoGAP control distinct local RhoA pools with different morphodynamic outputs (Fusco et al. 2016). Recent optogenetic work showed that DLC1 regulates Rho activity globally at steady state and locally at focal adhesions under tension, indicating that DLC1 output depends on subcellular location and mechanical state (Hinderling et al. 2026). Because the AHPH sensor reports active Rho at a relatively broad spatial scale, smaller DLC1-regulated RhoA pools at adhesions or protrusions may remain unresolved. Our data extend the emerging view of DLC1 as a spatial regulator of RhoA signaling, by showing that in 3D collagen invasion, DLC1 downregulation stabilizes rear-polarized RhoA activity and promotes multicellular streaming in a contractility-dependent manner.

Notably, beyond RhoA-driven contractility, our study reveals an additional, non-catalytic function of DLC1 in controlling how cancer cells engage with collagen-rich ECM. We discovered that DLC1 downregulation increases total and surface β1 integrin levels via a mechanism requiring the DLC1 LD-like motif. Functionally, β1 integrin blockade fully suppressed the invasive phenotype. Earlier work established the LD-like motif as a talin- and FAK-binding element required for focal adhesion localization and full tumor-suppressive DLC1 activity (Li et al. 2011; Zacharchenko et al. 2016). More recent work expanded this to a more dynamic model, in which force-dependent unfolding of the talin R8 domain reduces DLC1 binding and relieves local RhoA suppression (Haining et al. 2018). A recent study in cardiomyocytes further showed that DLC1–talin binding is tuned by the ECM stiffness and FAK signaling to regulate RhoA in a stiffness-dependent manner (Marhuenda et al. 2025). Together, these studies position the DLC1–talin interaction as a mechanosensitive module controlling adhesion-localized RhoA activity. The high affinity of DLC1 for the talin R8 domain has further been proposed to interfere with RIAM1 binding to talin, a process required for the membrane recruitment of talin and inside-out activation of integrins (Lee et al. 2009; Dahal et al. 2022). Our data support such a mechanism and provide evidence that the LD-like motif functions not only as a passive localization element, but also as an active modulator through which DLC1 limits β1 integrin enrichment and ECM adhesion during invasion in collagen-rich matrices.

However, the LD/talin mechanism likely explains only part of the β1 phenotype. As the shared β subunit of the major collagen-binding integrins, β1 integrin is frequently transcriptionally upregulated in tumors, where it has been linked to invasion, survival, therapy resistance, and metastatic progression (Chastney et al. 2025). Upon DLC1 depletion, β1 integrin protein and surface localization were increased without changes in mRNA levels, suggesting additional post-transcriptional regulation of β1 integrin. Besides altered trafficking or turnover, this could include post-translational mechanisms such as β1 integrin phosphorylation, which was recently implicated in breast cancer cell invasion (Conway et al. 2025). Previous work from our group showed that the related DLC family member DLC3, which localizes to endosomal membranes, regulates Rab4-dependent fast loop recycling of cargo proteins (Noll et al. 2019). Thus, DLC-family proteins may regulate invasion by controlling both adhesion-localized Rho signaling and the surface availability of receptors and proteases required for matrix invasion.

Together, our findings support a model in which DLC1 restrains 3D streaming invasion of cells by spatially limiting both β1 integrin engagement and RhoA–myosin contractility. Collagen cleavage may further promote path generation, but it cannot maintain streaming when myosin II activity is blocked. Thus, productive streaming requires β1 integrin–dependent adhesion and RhoA–myosin contractility to act together, allowing DLC1-low cells to convert collagen engagement into persistent multicellular invasion. Despite the molecular insights provided by this study, several questions remain. Future work should address how DLC1 controls β1 integrin adhesion-dependent stabilization, recycling and degradation depending on ECM composition and the associated α chain partner. Finally, while *ex vivo* cultures of patient-derived organoids support the conservation of the DLC1–β1 integrin–contractility axis, *in vivo* studies will be needed to test whether this module controls local invasion, collagen remodeling, dissemination or metastatic colonization in native tumor microenvironments.

## Material and Methods

### Cell lines and culture conditions

The following breast cancer cell lines were used in this study: MDA-MB-231 (obtained from CLS Cell Lines Service GmbH), BoM-1833 (kindly provided by Joan Massagué, Memorial Sloan Kettering Cancer Center, USA) and LentiX HEK293 (kindly provided by Philipp Rathert, Institute of Biochemistry, University of Stuttgart, Germany). Stable cell lines were cultured in media supplemented with the corresponding selection antibiotics: Blasticidin (8 µg/mL) and G418 (1000 µg/mL). MDA-MB-231 and BoM-1833 cells were maintained in DMEM supplemented with 10% fetal calf serum (FCS). LentiX HEK293 cells were maintained in DMEM medium with 10% FCS, 10 mM HEPES and 1 mM Sodium Pyruvate. All cell lines were cultured at 37 °C and 5% CO₂. All cell lines were authenticated, tested negative for mycoplasma (Lonza, LT07-318) and were kept in culture for no longer than 2 months.

### Organoid culture

Patient-derived breast cancer organoids were established from pleural effusion material obtained from an advanced breast cancer patient treated at the University Women’s Hospital, Tübingen (Önder et al. 2023). Use of human material was approved by the Ethics Committee of the Medical Faculty of the Eberhard Karls University Tübingen (150/2018BO2), compliant with all relevant ethical regulations regarding research involving human participants and informed consent was obtained. For full patient characteristics see Supplementary table S1. Organoids were cultured in advanced DMEM/F12 medium (1% Pen/Strep, 1x GlutaMAX^TM^, 10mM HEPES) supplemented with Neuregulin-1 (5 nM), FGF7 (5 ng/mL), FGF10 (20 ng/mL), A83-01 (500 nM), SB202190 (500 nM), Y-27632 (10 µM), B27 supplement (1×), N-acetylcysteine (1.25 mM) and 50% (v/v) L-WRN conditioned medium. L-WRN conditioned medium (ATCC #CRL-3276 (Miyoshi and Stappenbeck 2013), containing Wnt3a, R-spondin 3, and Noggin). Organoids were generated and maintained with minor adaptations of established protocols (Önder et al. 2023). In brief, dissociated cells were embedded as single cells in Matrigel at a ratio of 30% cell suspension to 70% Matrigel. Twenty-microlitre domes were plated into 48-well plates, allowed to polymerize for 30 min at 37 °C and 5% CO₂, and overlaid with organoid culture medium. Medium was refreshed every 3–4 days, and organoid growth was monitored by brightfield microscopy. Organoids were passaged every 5-9 days depending on size and density. For passaging, Matrigel domes were resuspended in ice-cold phosphate buffered saline (PBS) supplemented with 10µM Y-27632, pelleted by centrifugation (500 × g, 5 min) and enzymatically dissociated using 1 mL TrypLE Express for 5 min at 37 °C. For continued culture, the desired cell number was resuspended in culture medium, and re-embedded in fresh Matrigel at a ratio of 30% cell suspension and 70% Matrigel and cultured as described before.

### Plasmids and molecular cloning

All primer sequences are provided in supplementary Table S2. Epigenetic repression of DLC1 isoform 2 was performed using the CRISPRoff effector (DNMT3A–DNMT3L–dCas9–KRAB; originally described by (Nuñez et al. 2021) in a doxycycline-inducible format, together with constitutively expressed sgRNAs targeting the DLC1 isoform 2 promoter. In the published CRISPRoff-v2.1 plasmid (Addgene #167981), the effector is expressed from a constitutive CAG promoter. To generate a doxycycline-inducible lentiviral vector, the CRISPRoff-v2.1 insert was amplified as two overlapping PCR fragments using primers CRISPRoff-v2.1 Insert 1 and Insert 2, and assembled by Gibson Assembly into NheI/AvrII-digested pCW57-MCS1-P2A-MCS2 (Addgene #89180), thereby placing CRISPRoff expression under a Tet-responsive promoter. sgRNAs were cloned by oligonucleotide annealing and ligation into an Esp3I-digested pW212-lenti-spsgRNA-Esp3I-NLS-mScarlet vector (Addgene #170810). Two independent sgRNAs targeting distinct regions of the DLC1 promoter were tested: sgDLC1#1 (5’-GGAGAGGTGCGGCCATGTCC-3’) from a published CRISPRoff screen (Nuñez et al. 2021) and sgDLC1#2 (5’-ACGGCCCCAGAAAGAAAGCG-3’) designed with the Benchling sgRNA tool. A non-targeting LacZ sgRNA (5’-GTGCGAATACGCCCACGCGAT-3’, (Nuñez et al. 2021)) was used as a control.

For DLC1 overexpression studies, GFP-tagged constructs encoding wild-type DLC1, a catalytically inactive mutant (K714E), a talin-binding–deficient ΔLD mutant (Li et al. 2011), and a corresponding double mutant were used; a GFP-only vector served as control. The pEGFP-DLC1 construct has been described previously (Frey et al. 2022). ΔLD variants were generated by PCR-based backbone amplification and ligation. All GFP-based constructs were subcloned into the doxycycline-inducible lentiviral vector pCW57-MCS1-P2A-MCS2 (Addgene #89180) by restriction ligation cloning.

The Rho-GTP biosensor construct pEGFPC1-AHPH was a gift from Alpha Yap (University of Queensland) and was used as described previously (Piekny and Glotzer 2008; Priya et al. 2015; Gaston et al. 2021). All constructs were sequence-verified by Sanger sequencing prior to use.

### Antibodies

The following primary antibodies were used: mouse anti-DLC1 (BD Cat# 612021, RRID: AB_399416, 1:500 in WB), mouse anti-CD29 (BioLegend Cat# 303009, RRID:AB_314325, 1:400 in FACS), mouse anti-CD29 12G10 (Santa Cruz Biotechnology Cat# sc-59827, RRID:AB_782089, 1:100 in 2D and 3D IF), rat anti-CD29 AIIB2 (Sigma-Aldrich Cat# MABT409, RRID:AB_2893323, 1:100 in 2D and 3D IF), rabbit anti-Col1 3/4 fragment (ImmunoGlobe Cat# 0217-050, RRID:AB_2893279, 1:100 in 3D IF), mouse anti-CD29 (BD Biosciences Cat# 610467, RRID:AB_2128060, 1:1000 in WB), mouse anti-RhoA (Santa Cruz Cat# sc-418, RRID:AB_628218, 1:200 in WB), rabbit anti-RhoB (Cell Signaling Cat# 2098, RRID:AB_2179103; 1:500 in WB), rabbit anti-GFP (Cell Signaling Technology Cat# 2956, RRID: AB_1196615, 1:1000 in WB), rabbit anti-Talin (Proteintech Cat# 82856-4-RR, RRID:AB_3086557, 1:1000 in WB), rabbit anti-GAPDH (Sigma-Aldrich Cat# G9545, RRID: AB_796208, 1:5000 WB). Phalloidin labelling probes Alexa Fluor^TM^ 633 (Cat#A22284) and Alexa Fluor™ Plus 405 (Cat# A30104) were purchased from Thermo Fisher Scientific (1:1000 in IF). HRP-labelled secondary goat anti-mouse and anti-rabbit IgG antibodies were purchased from Dianova (Hamburg, Germany), Alexa-Fluor-labelled secondary IgG antibodies were from Invitrogen (1:1000 in FACS and 2D/3D IF). DAPI was from Sigma-Aldrich (5 µg/mL in IF).

### Lentiviral production and generation of stable cell lines

Lentiviral particles were produced in LentiX HEK293 cells by PEI-mediated transfection of the respective lentiviral expression vectors together with the packaging plasmid psPAX2 (Addgene #12260) and the envelope plasmid pCMV_VSV-G (Addgene #8454). Viral supernatants were collected 24 h post-transfection, filtered and used for transduction of target cells in the presence of polybrene (8 µg/mL). Stable MDA-MB-231 CRISPRoff cell lines were generated by co-transduction with equal volumes of lentiviral supernatants encoding the doxycycline-inducible CRISPRoff effector (pCW57-CRISPRoff-v2.1) and the respective sgRNA constructs (1:1 ratio of viral supernatants). Where indicated, CRISPRoff/sgRNA-expressing cell lines were subsequently re-transduced with a lentivirus encoding the GFP-AHPH Rho-GTP biosensor. For generation of stable CRISPRoff-expressing organoids, established organoid cultures were dissociated into single cells as described above and transduced in low-attachment 96-well plates with equal volumes of lentiviral supernatants encoding the CRISPRoff effector and sgRNA constructs (1:1 ratio). The same lentiviral supernatants used for 2D cell line transductions were used for transduction of organoid cultures. For this purpose, ttransductions were performed for 4 h at 37 °C in the presence of polybrene (8 µg/mL) and Y-27632 (10 µM). Following transduction, organoids were washed extensively with PBS, re-embedded in 70% Matrigel and returned to standard organoid culture conditions. Stable BoM-1833 cell lines were generated by single lentiviral transduction with doxycycline-inducible eGFP-tagged DLC1 constructs or an eGFP-only control.

Antibiotic selection was initiated 48 h post-transduction for both 2D cell lines and organoid cultures. 2D cell lines were selected with G418 (1000 µg/mL) and Blasticidin (8 µg/mL), whereas organoid cultures were selected with G418 (100 µg/mL) and Blasticidin (1 µg/mL) for 14 days. Transgene expression was induced by addition of 2 μg/mL doxycycline (Sigma).

### RNA interference and plasmid transfection

For transient knockdowns, cells were reverse transfected using Lipofectamine® RNAiMAX (Invitrogen) according to the manufacturer’s instructions. Target sequences for siRNAs are listed in supplementary Table S3. Cells were transfected with siRNAs at a final concentration of 5 nM and analysed 72 h post-transfection. siRNAs used in this study: siRhoA (ON-TARGETplus SMARTpool human RhoA L-003860-00-0005), siRhoB (ON-TARGETplus SMARTpool human RhoB L-008395-00-0005) from Thermo Scientific. Negative control siNT (ON-TARGETplus® non-targeting control pool D-001810-10 from Dharmacon).

### Spheroid culture and invasion assays

3D spheroid formation and invasion assays were performed in low-attachment, U-bottom 96-well plates (Nunclon™ Sphera™). MDA-MB-231 cells with CRISPRoff/sgRNA systems were treated with doxycycline (2 µg/mL) 24 h prior to spheroid formation and throughout the experiment, whereas BoM-1833 cells with DLC1 overexpression were induced only upon embedding of pre-formed spheroids into collagen. For spheroid generation, cells were resuspended at 1 × 10⁴ cells/mL in ice-cold complete medium supplemented with 2.5% (v/v) Matrigel and, where indicated, doxycycline and selection antibiotics. Aliquots of 100 µL (1,000 cells) were dispensed into pre-chilled plates, centrifuged (125 × g, 10 min, 8 °C), and incubated at 37 °C for 48 h to allow spheroid formation. Spheroids were then embedded in a collagen I matrix. This was prepared by neutralizing FibriCol collagen I (10 mg/mL) with 10× PBS and 0.1 M NaOH (and 0.1 M HCl if required) and diluting with 1× PBS to a final concentration of 1 mg/mL (pH 7.0–7.4). 100 µL of the neutralized collagen was added per well, plates were centrifuged briefly (300 × g, 3 min, 4 °C), and collagen was polymerized for 90 min at 37 °C. Complete medium (50 µL) containing doxycycline and/or inhibitors was then overlaid and refreshed every 2–3 days. Inhibitors were used at the following concentration: Blebbistatin (50 µM), GM6001 (50 µM), AIIB2 (1 µg/mL). Spheroid invasion was monitored by time-lapse brightfield imaging using the IncuCyte® S3 (4× objective, 1 h intervals for 5 days).

### Organoid collagen invasion assays

Organoid invasion assays were performed as described for spheroid invasion assays with the following modifications. The epigenetically edited organoids were cultured in media supplemented with 2 µg/ml doxycycline for a total of six days prior to invasion assays, with Y-27632 removed from the culture medium at the start of doxycycline addition and excluded throughout baseline invasion conditions. Flat-bottom 96-well plates were pre-coated with 30 µl Matrigel per well and polymerized at 37 °C. Organoids were harvested, dissociated into single cells and resuspended in ice-cold organoid medium. Cell suspensions were adjusted to 4 × 10⁴ cells/mL and supplemented with 2.5% (v/v) Matrigel. Collagen I matrices were prepared and neutralized as described above, and equal volumes of cell suspension and collagen solution were mixed to obtain a final collagen concentration of 1 mg/mL and 4,000 cells per well. Cell–collagen mixtures were dispensed into the pre-coated plates, centrifuged briefly (125 × g, 3 min, 4 °C), and incubated at 37 °C for 90 min to allow collagen polymerization. Complete organoid culture medium containing doxycycline (2 µg/mL) and inhibitors, where indicated, was gently overlaid and refreshed every 2–3 days. Inhibitors were used at the following concentration and added after polymerization: Y-27632 (50 µM) and AIIB2 (1 µg/mL). Organoid invasion was monitored for up to five days by time-lapse brightfield imaging using the IncuCyte® system (10× objective). Organoid growth was quantified using the IncuCyte® analysis software by measuring phase-contrast confluency and mScarlet fluorescence intensity over time, normalized to day 0 for each condition. Only mScarlet-positive organoids were included in the analysis.

### Dissociation of spheroids and organoids for flow cytometry or immunoblotting analyses

For downstream analysis, collagen-embedded spheroids and organoids were dissociated into single-cell suspensions. For each condition, spheroids (∼12–24 spheroids per sample) or organoids (pooled from 8–12 wells per condition) were collected, transferred to low-adhesion microcentrifuge tubes and briefly centrifuged (500 × g, 3 min) to remove residual medium. Collagen matrices were digested by incubation with Collagenase B (1.25 mg/mL in HBSS) at 37 °C with agitation until complete matrix dissolution (5–15 min). Samples were gently mixed during incubation to ensure efficient digestion. The reaction was stopped by addition of EDTA (0.5 mM), followed by centrifugation and the supernatant was discarded. The collagen-free material was subsequently dissociated into single cells by incubation at 37 °C for 5–10 min with either 1× Trypsin (spheroids) or TrypLE (organoids), with gentle mechanical trituration using low-adhesion pipette tips. Cells were collected by centrifugation (500 × g, 3 min) for downstream analyses.

### RNA isolation and RT-qPCR

RNA was extracted using the NucleoSpin® RNA Kit (Macherey-Nagel) according to the manufacturers’ instructions. RT–qPCR was performed using 100 ng total RNA per sample with a one-step, SYBR Green–based assay (Power SYBR® Green RNA-to-CT™ 1-Step Kit, Applied Biosystems) on a CFX96 Touch™ Real-Time PCR Detection System (Bio-Rad). Reactions were run in technical triplicates, and relative gene expression was calculated using the ΔCt method and normalized to the housekeeping gene RPLP0. qPCR primers used in this work were: RPLP0 Fw: 5’- CTCTGCATTCTCGCTTCCTGGAG-3’; RPLP0 RP: 5’- CAGATGGATCAGCCAAGAAGG-3’; DLC1 Fw: 5’- TTCCCCACAGCGCTTCCG-3’; DLC1 RP: 5’-TCGATGGGGAACAGGAAATCTTCA-3’; ITGB1 Fw: 5’-GGATTCTCCAGAAGGTGGTTTCG-3’; ITGB1 Rw: 5’-TGCCACCAAGTTTCCCATCTCC-3’.

### Cell lysis, SDS-PAGE and western blotting

Western blotting was performed on both 2D monolayer cultures and 3D collagen-grown cultures. For 2D cultures, the cells were plated on 10 µg/mL Collagen-R (Serva) coated dishes. Cells were washed with ice-cold PBS and lysed on ice in RIPA buffer composed of 50 mM Tris–HCl (pH 7.5), 0.1% (w/v) SDS, 150 mM NaCl, 0.25% sodium deoxycholate, 1% NP40, 1 mM sodium orthovanadate, 10 mM sodium fluoride, 20 mM β-glycerophosphate and 0.5 mM PMSF, supplemented with cOmplete™ EDTA-free protease inhibitor cocktail (Roche). Lysis was carried out for 30 min on ice, followed by clarification of lysates by centrifugation at 16,000 × g for 10 min at 4 °C. Protein concentrations were determined using the Bio-Rad DC Protein Assay according to the manufacturer’s instructions. Equal amounts of protein were resolved by SDS–PAGE on 4–12% Bis–Tris gels (NuPAGE® Novex®, Invitrogen) and transferred to nitrocellulose membranes using an iBlot® transfer system (Invitrogen). Membranes were blocked for 1 h at room temperature in blocking solution containing 0.5% Roche blocking reagent in PBS supplemented with 0.05% Tween-20 and 0.01% thimerosal, followed by incubation with primary antibodies at 4 °C overnight and HRP-conjugated secondary antibodies for 1 h at room temperature. Antibodies were diluted in blocking solution. Chemiluminescent signals were detected using an Amersham™ Imager 600 or 800 system (GE Healthcare) and quantified from 16-bit images acquired within the linear range using ImageQuant software.

### GFP-trap

BoM-1833 cell lines expressing GFP-tagged constructs were seeded at 2 × 10⁶ cells per 10-cm dish and induced with doxycycline 24 h after seeding; cells were harvested for GFP-Trap pulldown after 48 h of induction. GFP-Trap beads were prepared using an established protocol (Katoh et al.). For each pulldown reaction, 18 ng of GST-coupled GFP nanobody, produced in-house from an expression construct obtained from Kazuhisa Nakayama (Addgene #61838), were incubated with Glutathione Sepharose beads (GE17-5132-01, Sigma) in GFP-Trap lysis buffer (50 mM Tris–HCl pH 7.5, 150 mM NaCl, 0.5% Triton X-100, 10% glycerol, 0.5 mM PMSF, 1 mM Na₃VO₄, 10 mM NaF, 20 mM β-glycerophosphate and EDTA-free protease inhibitor cocktail) for 1 h at 4 °C with end-over-end rotation. Beads were washed twice with ice-cold lysis buffer prior to use for affinity purification. Cells were then lysed in GFP-Trap lysis buffer, incubated on ice for 10 min and clarified by centrifugation at maximum speed for 10 min at 4 °C. Protein concentrations were determined and adjusted to equal concentrations across samples. For each condition, equal amounts of clarified lysate were incubated with GFP-Trap beads for 2 h at 4 °C with rotation. A fraction of each lysate was retained as input. Beads were washed four times in lysis buffer to remove unbound material and collected by centrifugation. Bound proteins were eluted by addition of LDS sample buffer and heating at 70 °C for 10 min. Samples were supplemented with DTT and analyzed by SDS–PAGE and immunoblotting as described above.

### Flow cytometry

For flow cytometry, cells were cultured on 10 µg/mL Collagen-R (Serva) or obtained from dissociated organoids, as described above. Cells were harvested, fixed at RT in 2% paraformaldehyde for 5 min, washed in FACS buffer (PBS, 5% FCS, 0.05% sodium azide). Cells were stained with mouse primary antibodies or matched isotype controls in FACS buffer for 1 h at 4 °C, followed by incubation with Alexa Fluor 488- or 647-conjugated anti-mouse secondary antibodies for 1 h at 4 °C. Unstained, secondary-only and single-color controls were included for gating and compensation. Flow cytometry was performed using a MACSQuant Analyzer (Miltenyi Biotec), and data were analysed with FlowJo v10.

### Imaging

Fluorescence imaging was performed on an LSM 980 Airyscan 2 confocal microscope (Carl Zeiss) or an AxioObserver microscope (Carl Zeiss) equipped with a CSU-X1 spinning disk module and a Photometrics Evolve 512 EMCCD camera. For each experiment, identical laser power, detector gain and acquisition settings were applied across conditions. Z-stacks were acquired as indicated and are shown as maximum-intensity or sum-slice projections. Linear brightness/contrast adjustments and quantification of mean fluorescence intensity (MFI) from regions of interest were performed in ZEN (Carl Zeiss) and/or Fiji. For all samples, the following excitation wavelengths and detection windows were used: 405 nm and 425-488 for DAPI and Alexa-Fluor®- 405 coupled probes; 488 nm and 495-550 nm for GFP; 569 nm and 573-621 nm for mScarlet and Alexa-Fluor®-546 coupled probes; 633 nm and 655-720 nm for phalloidin-633 and Alexa-Fluor®- 633 coupled probes.

### Live-cell imaging of collagen embedded spheroids

Spheroid formation, maturation and collagen embedding were performed as described above. For live-cell imaging, individual spheroids derived from MDA-MB-231 cells expressing the GFP-AHPH Rho-GTP biosensor were transferred prior to collagen polymerization into poly(2-hydroxyethyl methacrylate) (polyHEMA)-coated glass-bottom imaging plates (10-well format) to prevent adhesion. Plates were centrifuged (300 × g, 3 min, 4 °C) and collagen polymerization and medium addition were carried out as described above. Live cell imaging was performed 6 days after embedding on an AxioObserver spinning disk system (Carl Zeiss) equipped with a CSU-X1 module and Photometrics Evolve 512 EMCCD camera. To maintain pH stability during imaging, medium was supplemented with 25 mM HEPES. For F-actin visualization, SPY650-FastAct (Spirochrome) was added 1 h prior to imaging. Time-lapse Z-stacks were acquired at 37 °C using a 20× objective with 15-min intervals (16 timepoints), capturing 1–2 positions per spheroid (maximum six positions per experiment). Image processing for display (maximum-intensity projections and linear brightness/contrast adjustments) was performed in ZEN.

### Immunofluorescence

For two-dimensional immunofluorescence analyses, cells were cultured on collagen-coated glass coverslips. Coverslips were coated with 10 µg/mL Collagen-R (Serva) for 2 h at 37 °C and washed with culture medium prior to seeding. MDA-MB-231 CRISPRoff cell lines were induced with doxycycline (2 µg/mL) for a total of 6 days and re-seeded onto coverslips (5 × 10⁴ cells per coverslip) on day 5 prior to fixation the following day. For experiments involving the GFP-AHPH Rho-GTP biosensor, MDA-MB-231 CRISPRoff/AHPH cells were induced with doxycycline for 6 days in total; for serum starvation assays, cells were pre-induced for 4 days prior to re-seeding. BoM-1833 cells expressing GFP-tagged DLC1 constructs were induced with doxycycline for 24 h, re-seeded onto coverslips and fixed after an additional 24 h. For knockdown experiments, transiently transfected cells were seeded at 5 × 10⁴ cells per coverslip 72 h post-transfection and fixed after 24 h. For serum starvation experiments, cells were washed 24 h after seeding and cultured in serum-free medium for an additional 24 h; where indicated, serum-containing medium was re-added for 10 min at 37 °C immediately prior to fixation. Cells were fixed and permeabilized simultaneously using 4% (w/v) paraformaldehyde and 0.1% (v/v) Triton X-100 in PBS for 10 min at room temperature. After blocking with 5% (v/v) goat serum for 1 h, cells were incubated with primary antibodies overnight at 4 °C. Following washing, fluorophore-conjugated secondary antibodies were applied for 1 h at room temperature; DAPI and/or phalloidin were included where indicated. Coverslips were mounted using ProLong™ Gold Antifade Mountant. Imaging was performed at RT using an LSM 980 Airyscan 2 (Carl Zeiss, Oberkochen, Germany) equipped with a Plan-Apochromat 63x/1.40 DIC or 40X/1.40 DIC (Carl Zeiss) oil immersion objective.

For the spheroid invasion assays, collagen-embedded spheroids or organoids were fixed with 4% paraformaldehyde for 40 min at room temperature and washed with PBS containing Ca²⁺ and Mg²⁺. For immunofluorescence staining, collagen plugs were permeabilized with 0.5% Triton X-100 in PBS for 30 min at room temperature, washed extensively with PBS, and blocked in 1% BSA in PBS for at least 1 h. Samples were incubated with specific primary antibodies diluted in blocking buffer overnight at 4 °C, followed by incubation with AlexaFluor® (488, 546, 633) labelled secondary antibodies and DAPI in blocking buffer overnight at 4 °C. After washing, samples were stored in PBS containing 0.02% sodium azide. Prior to microscopy, collagen plugs were transferred to 96-well glass-bottom imaging plates and imaged using a spinning disk confocal microscope equipped with a 63× oil-immersion objective. Images were acquired at RT using a AxioObserver microscope (Carl Zeiss) equipped with a CSU-X1 spinning disk module, a Photometrix Evolve 512 EMCCD camera equipped with a Plan-Apochromat 63x/1.40 DIC (Carl Zeiss) oil immersion objective or Plan-Apochromat 10x/0.45 M27 air objective.

### Image analysis and quantification IncuCyte image processing

Image analysis for images acquired with the IncuCyte® S3 was performed on phase-contrast or brightfield images exported as raw 8-bit files using Fiji (ImageJ). For each condition, approximately 10-25 spheroids per condition were analyzed.

### Spheroid invasive area

Spheroid outgrowth and invasion area was quantified from brightfield images by manually outlining the spheroid core and the total spheroid area using Fiji; invasive area was calculated as the difference between total and core area and all values were normalized to the mean of the corresponding experiment. Time-lapse image sequences were aligned using the Fiji plugin “Image Stabiliser”.

### Organoid morphology and protrusion analysis

Organoid morphology was assessed from phase-contrast images by exporting cell masks generated with the IncuCyte® analysis tool. Organoids > 2500 µm^2^ and < 300 µm^2^ were excluded from analysis. Shape descriptors, including circularity, solidity, and aspect ratio, were extracted using Fiji. A composite protrusion score was calculated from the shape descriptors and organoids were classified as compact or protrusive based on this score. The proportion of organoids in each category was quantified per condition.

### Confocal image processing

Image analysis for confocal microscopy was performed on raw CZI files using Fiji (ImageJ) with the Bio-Formats importer. All analyses were conducted on sum-slice projections unless stated otherwise. For each condition, approximately 20–25 cells were analysed in 2D experiments and 2–8 spheroids per condition were analysed in 3D experiments, corresponding to ∼20–80 cells per spheroid.

### Spatial distribution of invading cells

For quantification of the spatial distribution of invading cells, maximum intensity projections (MIPs) were generated from confocal image stacks encompassing entire spheroids. Individual nuclei were segmented based on the nuclear-localised mScarlet signal by thresholding in Fiji. The spheroid core centre and core boundary were defined, and the distance of each invading cell from the spheroid core was calculated. Invading cells were assigned to three distance-based zones (<400 µm, 400–550 µm and >550 μm from the spheroid core). The proportion of cells within each zone was calculated relative to the total number of invading cells per spheroid.

### β1 integrin and F-actin fluorescence analysis

For quantification of CD29 and actin signal distribution in 3D, cell masks were generated based on phalloidin staining using intensity thresholding in Fiji. From these masks, regions corresponding to the total cell area, plasma membrane and intracellular compartment were defined. Mean fluorescence intensity (MFI) was measured from sum-slice projections comprising five optical sections (two sections above and below the cell center). Plasma membrane enrichment was assessed by calculating the ratio of plasma membrane to intracellular CD29 signal. Whole-cell fluorescence intensities were quantified from sum-slice projections generated using a fixed number of optical sections per cell. The same number of slices was used for all cells within an experiment. Whole-cell fluorescence intensities were normalized to the mean value of the corresponding control condition within the same experiment.

### Collagen I cleavage analysis

Collagen I cleavage was quantified from 3D image stacks following pre-processing of the collagen and actin channels, including background subtraction and 3D median filtering (rolling background radius = 50; median filter: x = 1, y = 1, z = 0). Thresholded binary masks were generated using fixed thresholds applied consistently across all samples within an experiment. Cleavage volume was calculated by counting positive voxels within the thresholded mask and multiplying by the voxel volume derived from image metadata. Collagen cleavage index was calculated as the ratio of collagen cleavage volume to actin volume.

### Spatial analysis of GFP-AHPH signal

For analysis of GFP-AHPH in invading cells in live-cell imaging experiments, ROIs were exported at defined time points (0, 0.5, 1, and 1.5 h), cropped and rotated to standardise cell orientation. Cells were segmented into rear, middle and front regions using a custom Python-based analysis pipeline and mean fluorescence intensity (MFI) was quantified for each region. The fluorescence intensities of the ROIs were normalised to the mean whole-cell MFI at the corresponding time point and further normalised to the mean of the control condition within the same experiment.

### Temporal variability of Rho-GFP polarity

Temporal variability in Rho-GTP polarity was quantified for individual cells using the coefficient of variation (CV). For each cell, the polarity ratio was calculated at 0, 0.5, 1 and 1.5 h by dividing the normalized GFP–AHPH intensity at the cell front by that at the cell rear. For each cell, the mean polarity ratio across the four time points was calculated. Temporal variability was then quantified as the coefficient of variation (CV), defined as the standard deviation of the four front/rear polarity-ratio values divided by their mean. Thus, lower CV values indicate a more stable polarity ratio over time, whereas higher CV values indicate greater temporal variability in RhoA-GTP polarity.

## Statistical analysis

All experiments were performed independently at least three times unless otherwise stated. No data were excluded from the analyses. Statistical analyses were performed using GraphPad Prism 10 (GraphPad Software). For analyses based on individual cells or organoids, measurements were summarized as replicate means for statistical inference unless otherwise indicated in the corresponding figure legends. Statistical inference was based on independent biological replicates.

Comparisons between two groups were performed using unpaired two-tailed Student’s t-tests, unpaired two-tailed t-tests with Welch’s correction, or two-tailed Mann–Whitney U-tests, as specified in the corresponding figure legends. Comparisons involving more than two groups were performed using one-way ANOVA followed by Tukey’s multiple-comparisons test. Experiments involving two independent factors were analyzed using ordinary two-way ANOVA, followed by Tukey’s or Šídák’s multiple-comparisons test, as specified in the corresponding figure legends. Distance-distribution data were analyzed using two-way repeated-measures ANOVA followed by Tukey’s multiple-comparisons test comparing sgRNA conditions within each distance category. No inferential statistical testing was performed for experiments with (n = 1).

For standard scatter and bar plots, individual points represent independent biological replicates and data are shown as mean ± s.d. unless otherwise stated. Beeswarm SuperPlots display individual measurements as small light-colored dots. Larger black dots represent the mean of each biological replicate, and error bars indicate the mean ± s.d. of the biological replicate means. Where individual cells, spheroids or organoids are shown, the plotted and statistical units are specified in the corresponding figure legends. Sample sizes and the statistical tests used are specified in the corresponding figure legends. p-values are indicated as follows: *P* ≤ 0.05 (*), *P* ≤ 0.01 (**), *P* ≤ 0.001 (***), and *P* ≤ 0.0001 (****); n.s., not significant.

## Acknowledgements

The authors gratefully acknowledge the Technology Platform “Cellular Analytics” of the Stuttgart Research Center Systems Biology for support and assistance in this work. We thank the Rathert group for providing Lenti-X cells and the Massagué laboratory for providing the BoM-1833 cells. We thank Nikolai Herdrich for assistance with production of viral particles. We thank Camille Dantzer for support with organoid characterization, and Barbara Volz for providing information on the organoid models and culture. We thank Angelika Hausser and Andrew Clark for critical reading of the manuscript and the scientific discussions throughout the project. Figures 1A and 6B were created with BioRender.com.

## Author Contributions

Conceptualization, M.A.O., C.L. and F.K.; Methodology, F.K., F.M. and A.K.; Investigation, F.K., S.E., O.P., L.T., and C.L.; Writing—original draft, F.K., C.L. and M.A.O.; Writing—review & editing, F.K., C.L. and M.A.O.; Funding acquisition, C.L., and M.A.O.; Supervision, M.A.O., C.L. and F.K.; Project administration, M.A.O.

## Data availability

All data supporting the conclusions of this study are presented in the manuscript, supplementary materials, and the source data files. Furthermore, uncropped western blots performed in this study are included in this publication. All data and materials reported in this publication will be shared by the lead contact upon request.

## Supplementary Figure Captions

**Supplementary Figure 1: Validation of DLC1 depletion and image analysis workflows.**

(A, B) Quantitative PCR analysis of DLC1 isoform 2 mRNA levels in MDA-MB-231 cells expressing control (sgLacZ) or DLC1-targeting sgRNAs. RNA was isolated from cells grown under 2D culture conditions (A) or as spheroids embedded in collagen I (B), following doxycycline induction for 6 days. n = 4-5 independent experiments, data are presented as mean ± s.d.; one-way ANOVA followed by Tukey’s multiple-comparisons test.

(C) Representative brightfield images of doxycycline-induced MDA-MB-231 CRISPRoff spheroids expressing sgLacZ or DLC1-targeting sgRNAs (sgDLC1#1). The spheroids were embedded in 1 mg/mL collagen I, imaged on day 0 (day of collagen addition) and day 5. Boxes indicate the magnified regions of interest (ROIs). Scale bar = 400 µm.

(D) Representative magnified brightfield images from time-lapse imaging at t = 68, 70 and 72 h of doxycycline-induced MDA-MB-231 CRISPRoff spheroids expressing sgLacZ or DLC1-targeting sgRNAs. The spheroids were embedded in 1 mg/mL collagen I. White arrows indicate the same individual cells tracked across frames, illustrating their displacement over time. Scale bar = 400 µm.

(E) Representative brightfield images of a collagen-embedded spheroid at day 0 and day 5 after collagen addition, with the segmentation masks used for invasive area quantification in Fig. 1F. The spheroid core (yellow region of interest (ROI 2) and total spheroid area (yellow region of interest ROI 1) were manually delineated and invasive area was calculated by subtracting the spheroid core area from the total spheroid area. Scale bar = 400 µm.

(F) Representative confocal images of doxycycline-induced MDA-MB-231 CRISPRoff spheroids expressing sgLacZ or DLC1-targeting sgRNAs (sgDLC1#1), embedded in 1 mg/mL collagen I and imaged at day 5. Nuclei are visualized by NLS–mScarlet fluorescence (black). Scale bar = 400 µm.

(G) Quantification of the total number of invading cells from the spheroid core, based on confocal images shown in (Fig.S1F). n = 3 independent experiments, each analyzing ≥5 spheroids per condition; data are presented as mean ± s.d.; one-way ANOVA followed by Tukey’s multiple-comparisons test.

(H) Representative brightfield images illustrating spheroid formation in growth medium supplemented with 2.5% Matrigel, Images were taken at 0, 12 h, and 48 h for control and DLC1-targeting cell lines, demonstrating comparable spheroid assembly kinetics. Scale bar = 400 µm.

(I) Quantification of spheroid core area of doxycycline-induced MDA-MB-231 CRISPRoff spheroids expressing sgLacZ or DLC1-targeting sgRNAs 48h after spheroid formation. n = 7 independent experiments, each analyzing ≥5 spheroids per condition; data are presented as mean ± s.d.; one-way ANOVA followed by Tukey’s multiple-comparisons test.

(J) Quantification of relative spheroid core area as shown representatively as in (E) at day 5 following treatment with vehicle control or GM6001 (50 µM), showing no significant difference between conditions. n = 4 independent experiments, each analyzing ≥5 spheroids per condition; data are presented as mean ± s.d.; two-way ANOVA with sgRNA and treatment as factors, followed by Tukey’s multiple-comparisons test.

(K) Representative brightfield images of GM6001 (50 µM) or vehicle control treated, doxycycline-induced MDA-MB-231 CRISPRoff spheroids expressing sgLacZ or DLC1-targeting sgRNAs (sgDLC1#1). The spheroids where embedded in 1 mg/mL collagen I, imaged at day 0 (day of collagen addition) and day 5. Boxes indicate the magnified regions of interest (ROIs). Scale bar = 200 µm.

(L) Representative doxycycline-induced MDA-MB-231 CRISPRoff spheroids expressing sgLacZ or DLC1-targeting sgRNAs (sgDLC1#1). The spheroids were embedded in 1 mg/mL collagen I, immunolabelled for MMP-cleaved collagen I (Col1-3/4; green, inverted grayscale look-up table (LUT)) and F-actin (magenta) and imaged on day 5. NLS-mScarlet was used as a nuclear marker. Insets show magnified regions of interest (ROIs) of invading cell strands. Scale bar = 400 µm.

**Supplementary Figure 2: Actomyosin organization and validation of the GFP–AHPH biosensor in DLC1-depleted cells**

(A) Representative immunofluorescence images of doxycycline-induced MDA-MB-231 CRISPRoff spheroids expressing sgLacZ or DLC1-targeting sgRNAs. Spheroids were embedded in 1 mg/mL collagen and stained for F-actin (magenta, grayscale LUT) after 5 days of invasion. Shown are maximum-intensity projections (MIPs). Scale bar = 30 µm.

(B) Quantification of total cellular F-Actin fluorescence intensity in invading cells per invading cell based on images as representatively shown in (A). n = 4 independent experiments, each analyzing ≥25 cells per condition; data are presented as mean ± s.d.; one-way ANOVA followed by Tukey’s multiple-comparisons test.

(C) Representative brightfield images of blebbistatin (10 µM) or vehicle control treated, doxycycline-induced MDA-MB-231 CRISPRoff spheroids expressing sgLacZ or DLC1-targeting sgRNAs (sgDLC1#1). The spheroids were embedded in 1 mg/mL collagen I and were imaged at day 0 (day of collagen addition) and day 5. Boxes indicate the magnified regions of interest (ROIs). Scale bar = 400 µm.

(D) Representative magnified immunofluorescence images of MDA-MB-231 CRISPRoff spheroids expressing sgLacZ or DLC1-targeting sgRNAs treated with blebbistatin (10 µM) or vehicle control (DMSO). The spheroids were embedded in 1 mg/mL collagen I, immunolabelled for MMP-cleaved collagen I (Col1-3/4; green) and F-actin (magenta) after 5 days of invasion. mScarlet–NLS was used as a nuclear marker. Scale bar = 100 µm.

(E) Quantification of spheroid core area at day 5 in collagen following treatment with vehicle control or blebbistatin (10 µM). n = 4 independent experiments, each analyzing ≥5 spheroids per condition; data are presented as mean ± s.d.; two-way ANOVA with sgRNA and treatment as factors, followed by Tukey’s multiple-comparisons test.

(F) Immunoblot analysis of whole-cell lysates from CRISPRoff doxycycline-induced MDA-MB-231 cells expressing control (sgLacZ) or DLC1-targeting sgRNAs and independently expressing GFP-AHPH WT. Cell lysates were prepared 24 h post-seeding. Probed for GFP. GAPDH served as a loading control.

(G) Representative immunofluorescence images of MDA-MB-231 cells with stable expression of the GFP-AHPH WT (green) reporter. Images were taken following 72h of RhoA, RhoB and combined RhoA/B siRNA knockdown. Cells were fixed and stained for F-actin (magenta) and nuclei (turquoise). Scale bar = 10 µm.

(H) Immunoblot analysis of whole-cell lysates from MDA-MB-231 cells expressing RhoA activity biosensor GFP–AHPH-WT (green) with the indicated siRNAs. Membranes were probed for RhoA and RhoB. The RhoB and double RhoA/B knockdown lanes were non-adjacent on the same membrane and are shown together for presentation. GAPDH was used as loading control.

(I) Representative brightfield images of GFP–AHPH WT, doxycycline-induced MDA-MB-231 CRISPRoff spheroids expressing sgLacZ or DLC1-targeting sgRNAs. The spheroids were embedded in 1 mg/mL collagen I, imaged at day 0 (day of collagen addition) and day 5. Boxes indicate the magnified regions of interest (ROIs). Scale bar = 500 µm. ROI Scale bar = 100 µm.

(J) Quantification of invasive area per spheroid after 5 days in collagen I based on images as representatively shown in (J). Data are presented as mean ± s.d. from ≥10 spheroids per condition in one experiment.

**Supplementary Figure 3: Expression and localization controls for β1 integrin.**

(A) Representative image of the invasive front of doxycycline-induced MDA-MB-231 CRISPRoff spheroids expressing sgDLC1#2. The spheroid was embedded in 1 mg/mL collagen I and imaged on day 5. Boxes show magnified ROIs, with white arrows highlighting representative invasion-associated tracks within the collagen matrix. Scale bar spheroid = 100 µm; scale bar ROI = 50 µm.

(B) Representative immunofluorescence images of invading cells from AIIB2 (1 µg/mL) or vehicle control treated, doxycycline-induced MDA-MB-231 CRISPRoff spheroids expressing sgLacZ or DLC1-targeting sgRNAs (sgDLC1#1). The spheroids were embedded in 1 mg/mL collagen I, stained for total β1 integrin (green, grayscale LUT), cleaved Col1 ¾ (magenta) and F-actin (red), and imaged at day 5 following invasion. Nuclei are shown in blue. Shown are single optical sections. Invading cells from sgLacZ and DLC1-depleted spheroids are shown. Scale bar = 30 µm

(C) Quantification of plasma membrane-to-cytoplasmic β1 integrin fluorescence intensity ratio in invading cells following AIIB2 (1 µg/ml) treatment shown in (C). n = 3 independent experiments, each analyzing ≥ 40 cells per condition. Data presented as mean ± s.d.; one-way ANOVA followed by Tukey’s multiple-comparisons test.

(D) Representative brightfield images of AIIB2 (1 µg/ml) or vehicle control treated doxycycline-induced MDA-MB-231 CRISPRoff spheroids expressing sgLacZ or DLC1-targeting sgRNAs (sgDLC1#1). The spheroids were embedded in 1 mg/mL collagen I, imaged at day 5. Boxes indicate the magnified regions of interest (ROIs). Scale bar = 400 µm.

(E) Quantification of spheroid core area at day 5 in collagen following treatment with vehicle control or AIIB2 (1 µg/mL). n = 4 independent experiments, each analyzing ≥5 spheroids per condition; data are presented as mean ± s.d.; two-way ANOVA with sgRNA and treatment as factors, followed by Tukey’s multiple-comparisons test.

(F) Quantitative PCR analysis of *ITGB1* mRNA levels in MDA-MB-231 cells expressing control (sgLacZ) or DLC1-targeting sgRNAs. RNA was isolated from cells grown under 2D culture conditions following doxycycline induction for six days. n = 3 independent experiments, data are presented as mean ± s.d.; one-way ANOVA followed by Tukey’s multiple-comparisons test.

**Supplementary Figure 4: Characterization of BoM-1833 cells, DLC1 re-expression and inhibitor treatments.**

(A) Quantitative PCR analysis of *DLC1 isoform 2* mRNA levels in MDA-MB-231 and BoM-1833 cells. RNA was isolated from cells grown under 2D culture conditions. *RPLP0* was used as housekeeping gene. n = 3 independent experiments, data are presented as mean ± s.d.; Unpaired two-tailed t-test.

(B) Representative brightfield images of GM6001 (50 µM), Blebbistatin (10 µM), AIIB2 (1 µg/mL) or vehicle control treated MDA-MB-231 and BoM-1833 spheroids. The spheroids were embedded in 1 mg/mL collagen I and imaged after 5 days. Scale bar = 400 µm.

(C) Quantification of invasive area per spheroid after 5 days in collagen I following treatment with GM6001 (50 µM), Blebbistatin (10 µM), AIIB2 (1 µg/mL) or vehicle control, based on images representatively shown in (B). n = 3 independent experiments, each analyzing ≥2 spheroids per condition. Data presented as mean ± s.d.; two-way ANOVA with cell line and treatment as factors, followed by Šídák’s multiple-comparisons test.

(D) Immunoblot analysis of whole-cell lysates from BoM-1833 cells expressing GFP, GFP–DLC1c(WT), or GFP–DLC1 (K714E), following 24 h of doxycycline treatment. Probed for DLC1. GAPDH was used as loading control.

(E) Representative immunofluorescence images showing the localization in green of GFP, GFP–DLC1 (WT) and GFP-DLC1 (K714E) in 2D cultures, co-stained with vinculin (orange) to indicate focal adhesions. Scale bar = 5 µm.

(F) Immunoblot analysis of whole-cell lysates of BoM-1833 cells expressing GFP, GFP–DLC1 (WT), or GFP–DLC1 (K714E), following 3 days of doxycycline induction and culture on collagen-coated dishes. Probed for β1 integrin and DLC1. GAPDH was used as loading control.

(G) Densitometry shows relative levels of β1 integrin quantified on immunoblots representatively shown in (F) n = 5 independent experiments, data are presented as mean ± s.d.; one-way ANOVA followed by Tukey’s multiple-comparisons test.

**Supplementary Figure 5: β1 integrin expression in BoM-1833 cells expressing DLC1 separation-of-function mutants.**

(A) Immunoblot analysis of whole-cell lysates from BoM-1833 cells expressing GFP control, GFP–DLC1 (WT), or GFP–DLC1 (K714E/ΔLD) following 3 days of doxycycline induction and cultured on collagen-coated dishes. Probed for β1 integrin and DLC1. GAPDH was used as loading control.

(B) Densitometry shows relative levels of β1 integrin quantified on immunoblots representatively shown in (A) n = 5 independent experiments, Data are represented as mean ± s.d.; one-way ANOVA followed by Tukey’s multiple-comparisons test.

**Supplementary Figure 6: DLC1 regulates a conserved β1 integrin–dependent invasion program in patient-derived metastatic organoids.**

(A) Representative brightfield and fluorescence images of breast cancer organoids embedded in Matrigel at 7 days after transduction, expressing sgLacZ or DLC1-targeting sgRNAs. NLS-mScarlet (red) was used as a nuclear marker. Scale bar = 75 µm.

(B) Representative brightfield and fluorescence images of doxycycline-induced CRISPRoff organoids expressing sgLacZ or DLC1-targeting sgRNAs. The organoids were embedded in 1 mg/mL collagen I and imaged at day 0 (day of collagen addition) and day 5. Scale bar = 400 µm

(C) Relative mScarlet–NLS fluorescence intensity in doxycycline-induced CRISPRoff organoids expressing sgLacZ or DLC1-targeting sgRNAs, used as a proxy for organoid growth and normalized to t = 0. Organoids were embedded in 1 mg/mL collagen I and monitored by live-cell imaging for 5 days.

(D) Immunoblot analysis of whole-cell lysates from CRISPRoff organoids expressing control (sgLacZ) or DLC1-targeting sgRNAs, following 6 days of doxycycline induction, probed for β1 integrin. GAPDH was used as loading control.

(E) Densitometry shows relative levels of β1 integrin quantified on immunoblots representatively shown in (D). n = 3 independent experiments.; data are presented as mean ± s.d.; one-way ANOVA followed by Tukey’s multiple-comparisons test.

**Supplementary Figure 7: Uncropped Western Blots.** Merged JPEG images display the colorimetric molecular weight marker overlaid with the chemiluminescence signal. Rectangles indicate the regions of the membrane that were cropped in the TIFF files and shown in the annotated main figures.

(A) Uncropped Western Blot images for Figure 1B.

(B) Uncropped Western Blot images for Figure 3F.

(C) Uncropped Western Blot images for Figure 4A.

(D) Uncropped Western Blot images for Figure 5A.

(E) Uncropped Western Blot images for Supplementary Figure 2F.

(F) Uncropped Western Blot images for Supplementary Figure 2H.

(G) Uncropped Western Blot images for Supplementary Figure 4D.

(H) Uncropped Western Blot images for Supplementary Figure 4F.

(I) Uncropped Western Blot images for Supplementary Figure 5A.

(J) Uncropped Western Blot images for Supplementary Figure 6D.

**Supplementary Movies 1:**

Time-lapse brightfield imaging of DLC1-dependent spheroid invasion in collagen. Doxycycline-induced MDA-MB-231 CRISPRoff spheroids expressing sgLacZ or DLC1-targeting sgRNAs were embedded in 1 mg/mL collagen I and imaged by brightfield microscopy every 1 h for 96 h. Scale bar = 400 µm.

**Supplementary Movie 2:**

Time-lapse brightfield imaging of patient-derived organoid invasion in collagen. Doxycycline-induced patient-derived CRISPRoff organoids expressing sgLacZ or DLC1-targeting sgRNAs were embedded in 1 mg/mL collagen I and imaged by brightfield microscopy every 1 h for 5 days after embedding. Representatively are shown movies from day 3 (t = 48) to day 5 ( t= 120). The video corresponds to the organoid invasion experiment as representatively shown in Fig. 6D. Scale bar = 100 µm.

**Supplementary Table S1:** List of BC patient data from whom the organoid line (#6) was established (Önder et al. 2023). Abbreviations: NST (no special type).

**Supplementary Table S2**: List of oligonucleotides used for molecular cloning.

**Supplementary Table S3**: siRNA sequences used for RNA interference.

