## Supplementary Figures for "DLC1 loss drives multicellular streaming invasion by enforcing spatially coordinated Rho and β1 integrin signaling"

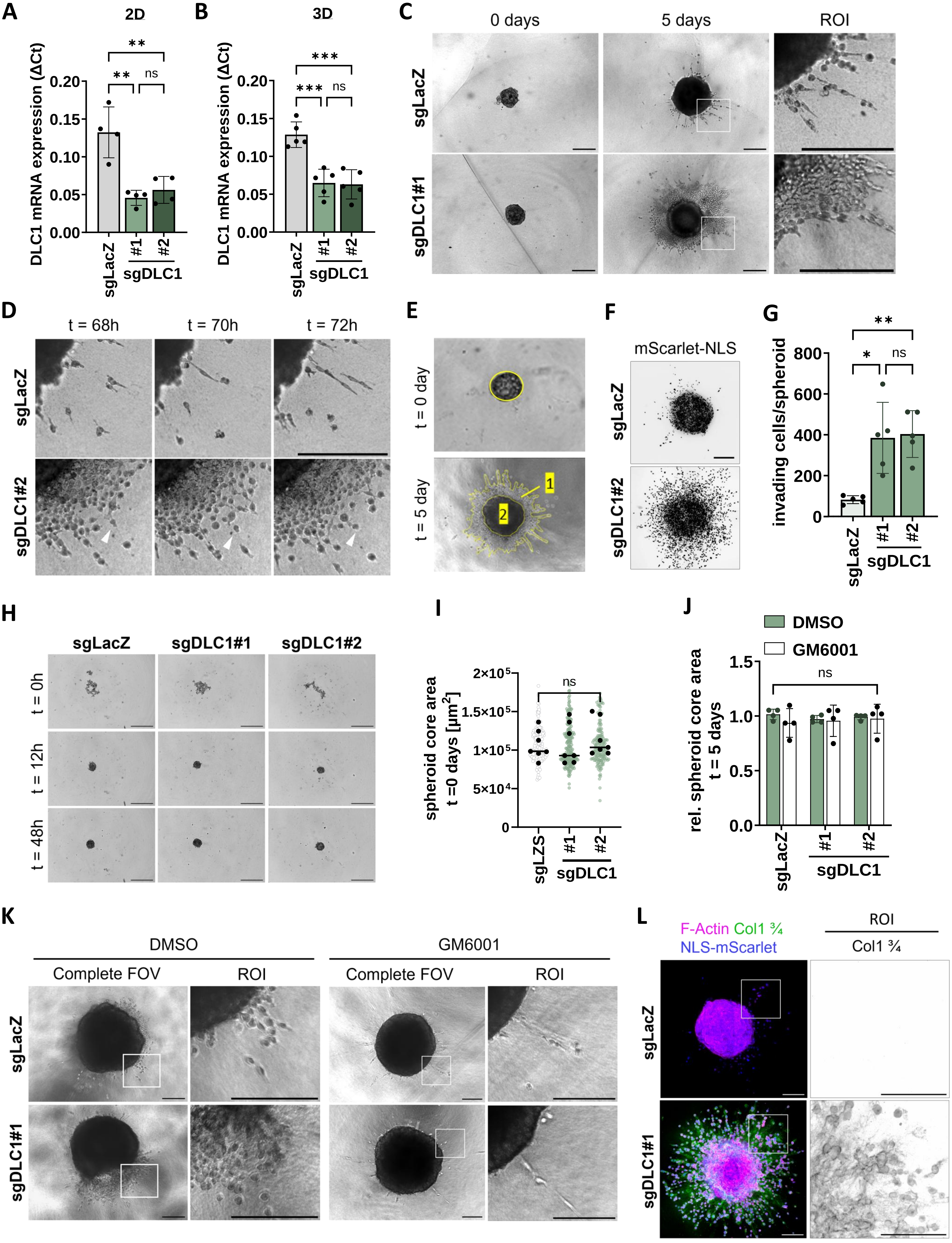


Supplementary figure 1

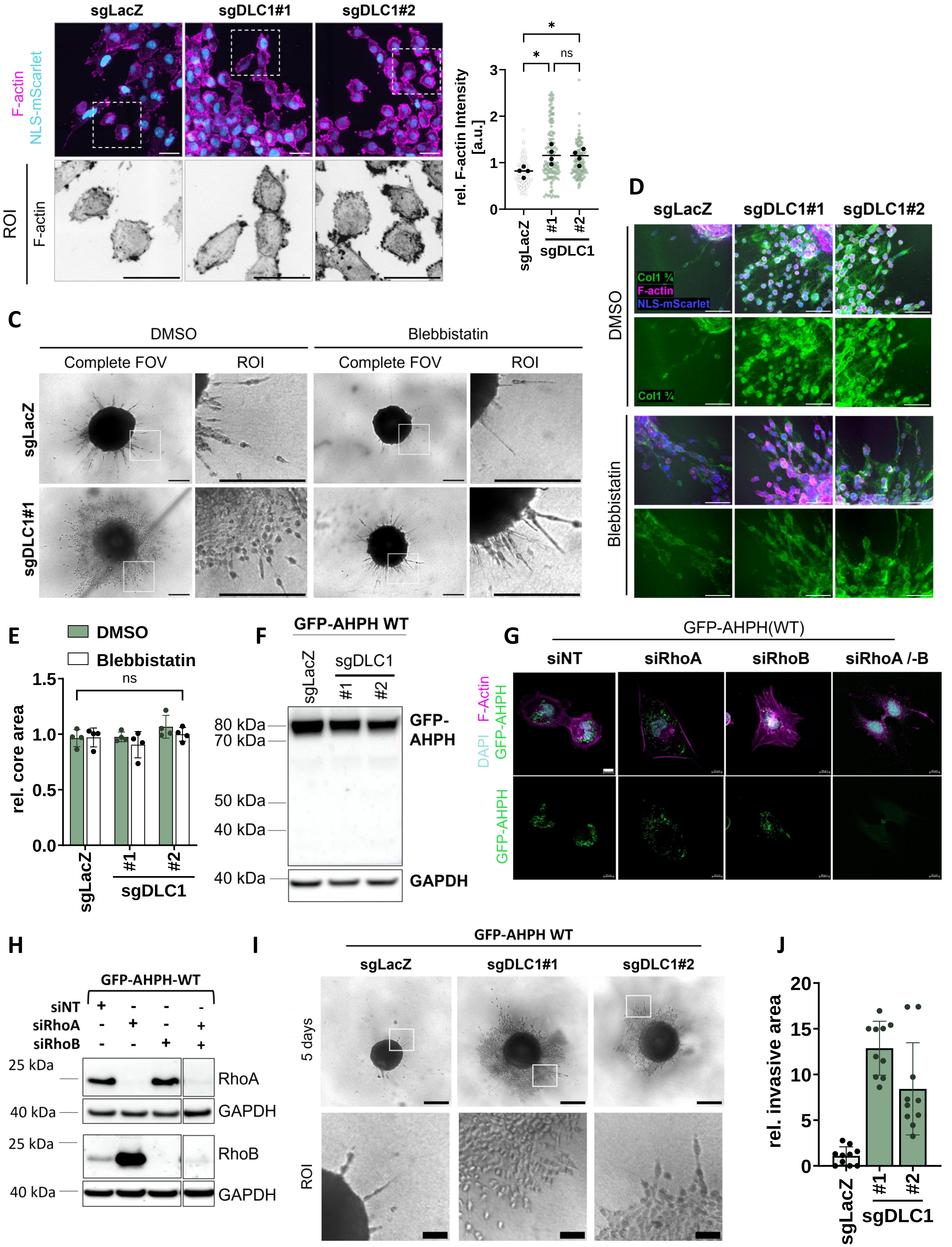
A
sgLacZ
sgDLC1#1
sgDLC1#2
F-actin
NLS-mScarlet
ROI
F-actin
B
rel. F-actin Intensity [a.u.]
sgLacZ
#1
#2
sgDLC1
ns
C
DMSO
Blebbistatin
Complete FOV
ROI
sgLacZ
sgDLC1#1
D
Col1 ¾
F-actin
NLS-mScarlet
E
rel. core area
F
GFP-AHPH WT
80 kDa
70 kDa
50 kDa
40 kDa
GFP-AHPH
GAPDH
G
GFP-AHPH(WT)
siNT
siRhoA
siRhoB
siRhoA /-B
DAPI
GFP-AHPH
F-Actin
H
GFP-AHPH-WT
25 kDa
RhoA
RhoB
I
5 days
J
rel. invasive area

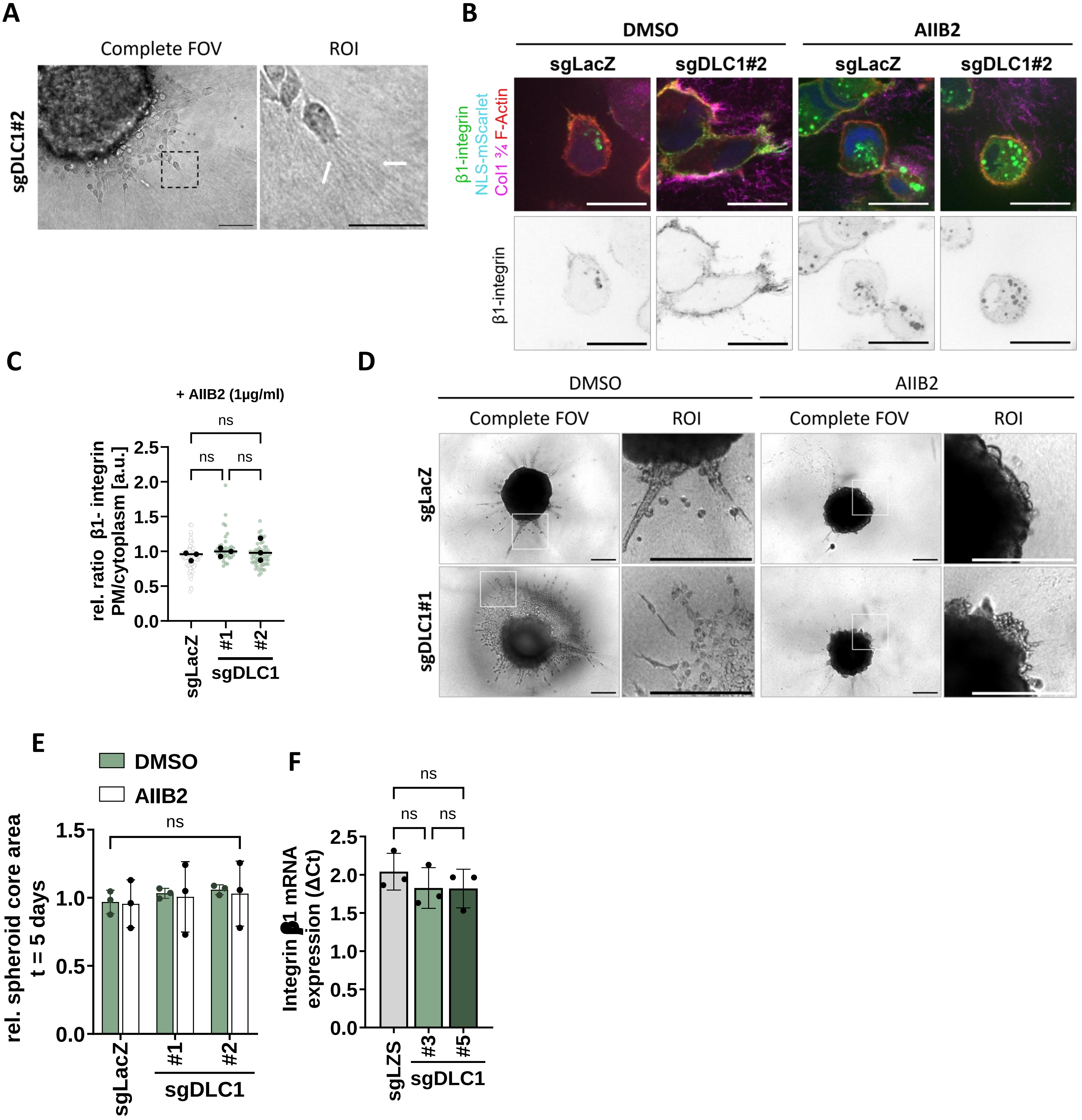


Supplementary figure 3

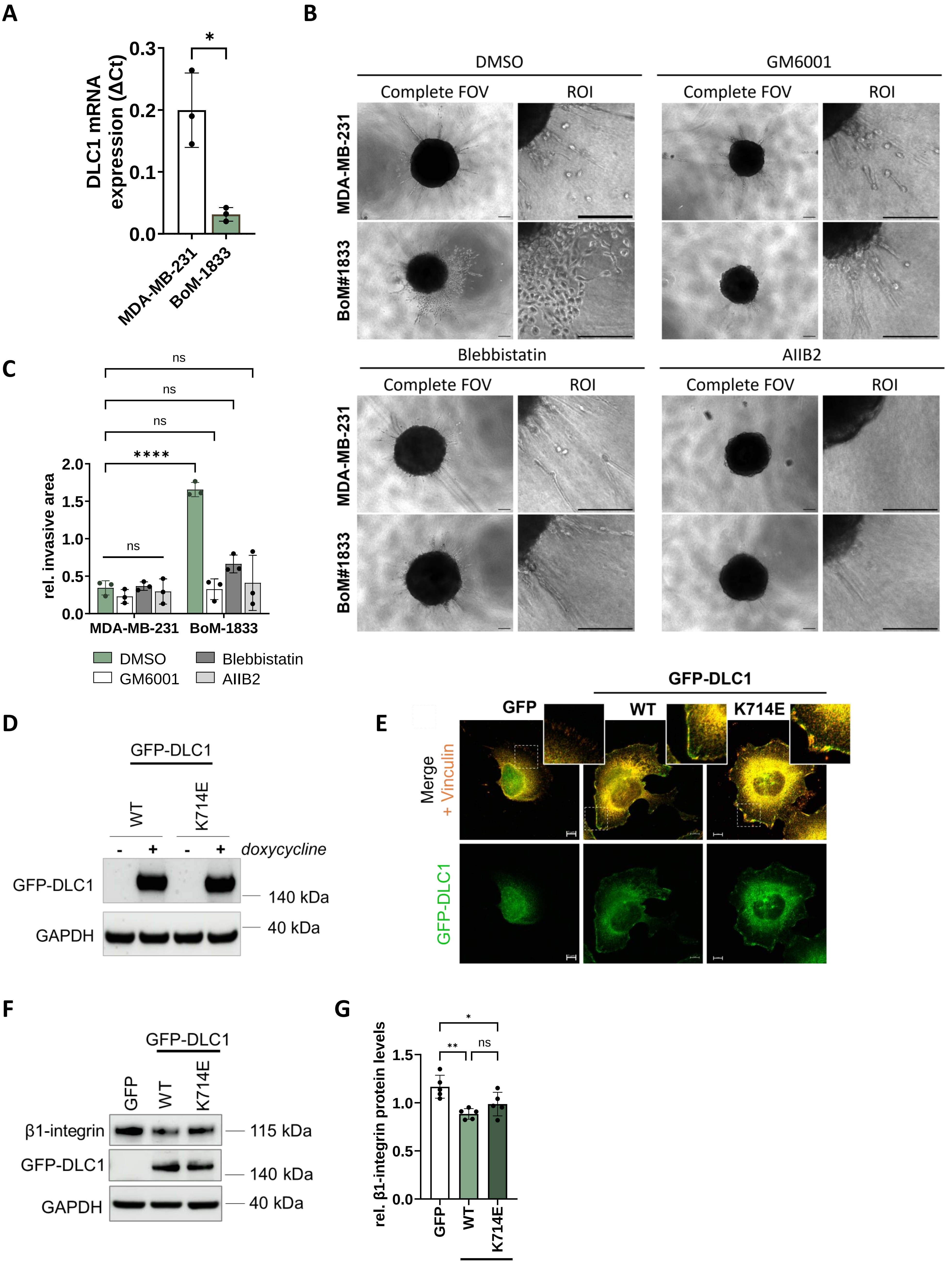


**Supplementary figure 4**

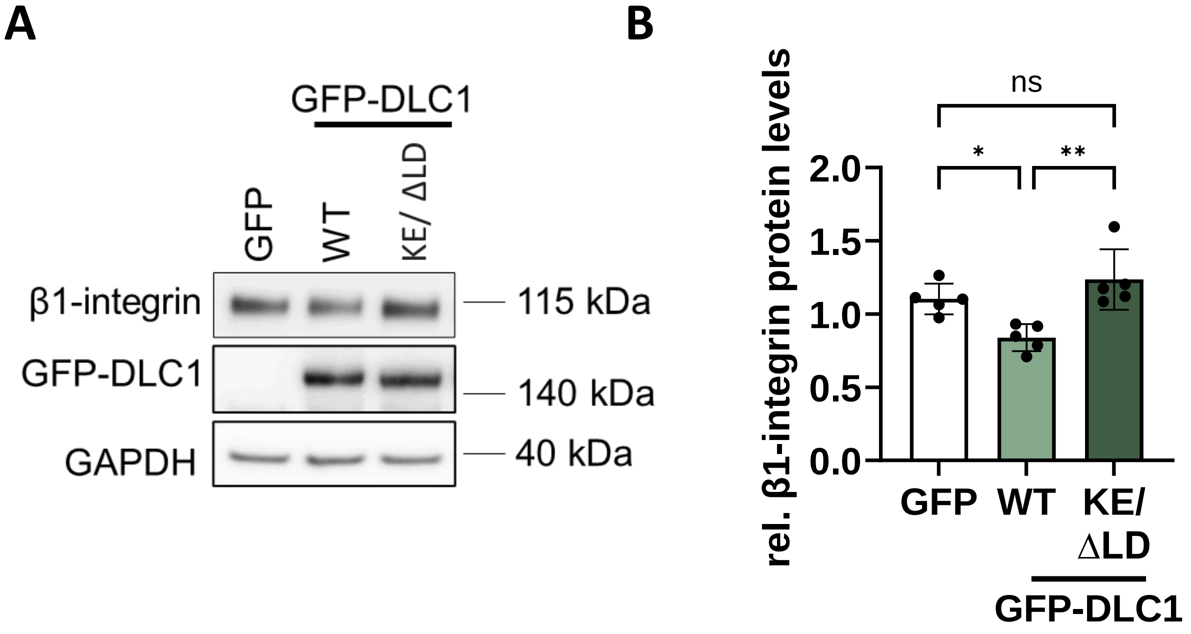


<u>**Supplementary figure 5**</u>

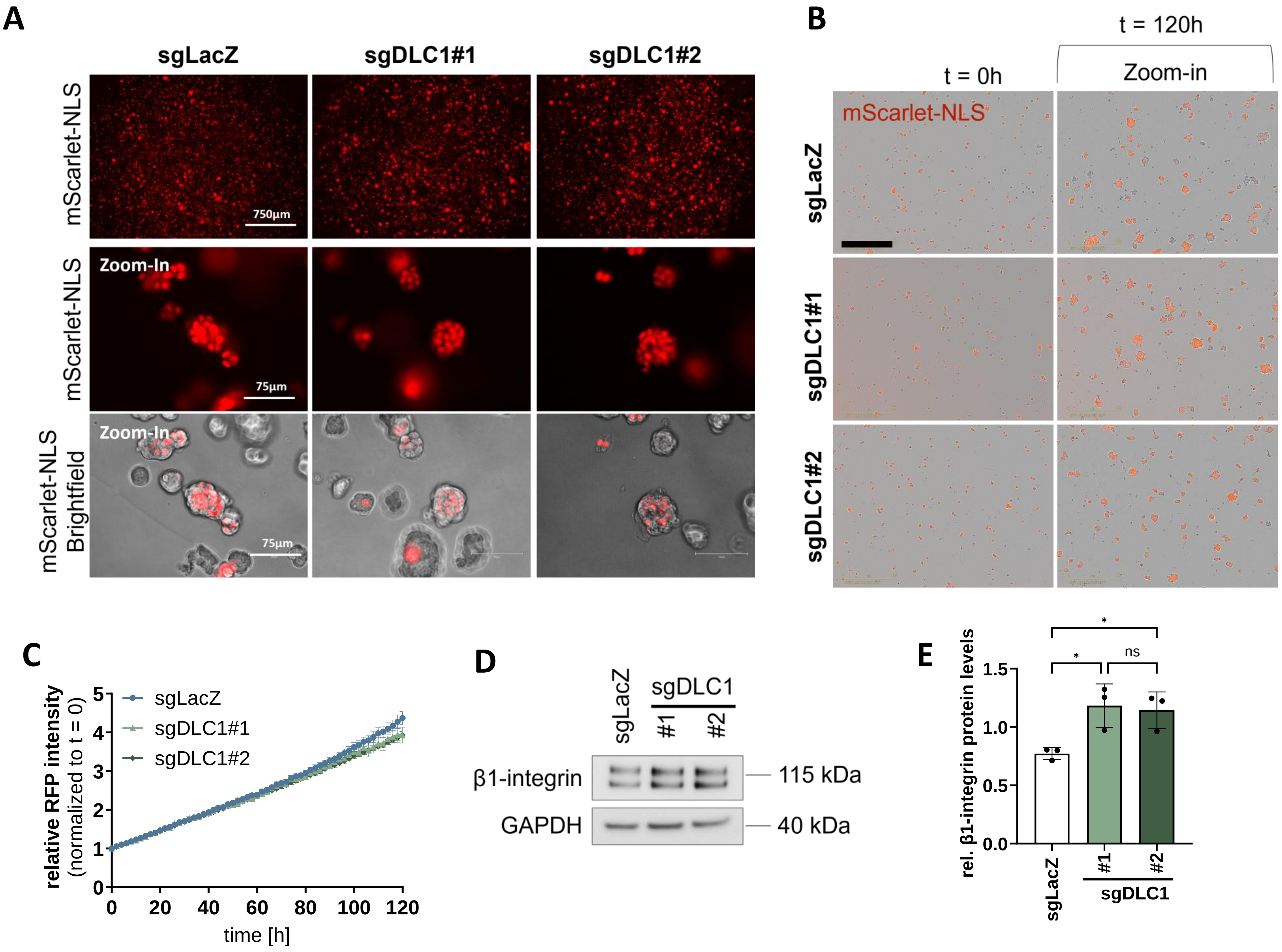


**Supplementary figure 6**

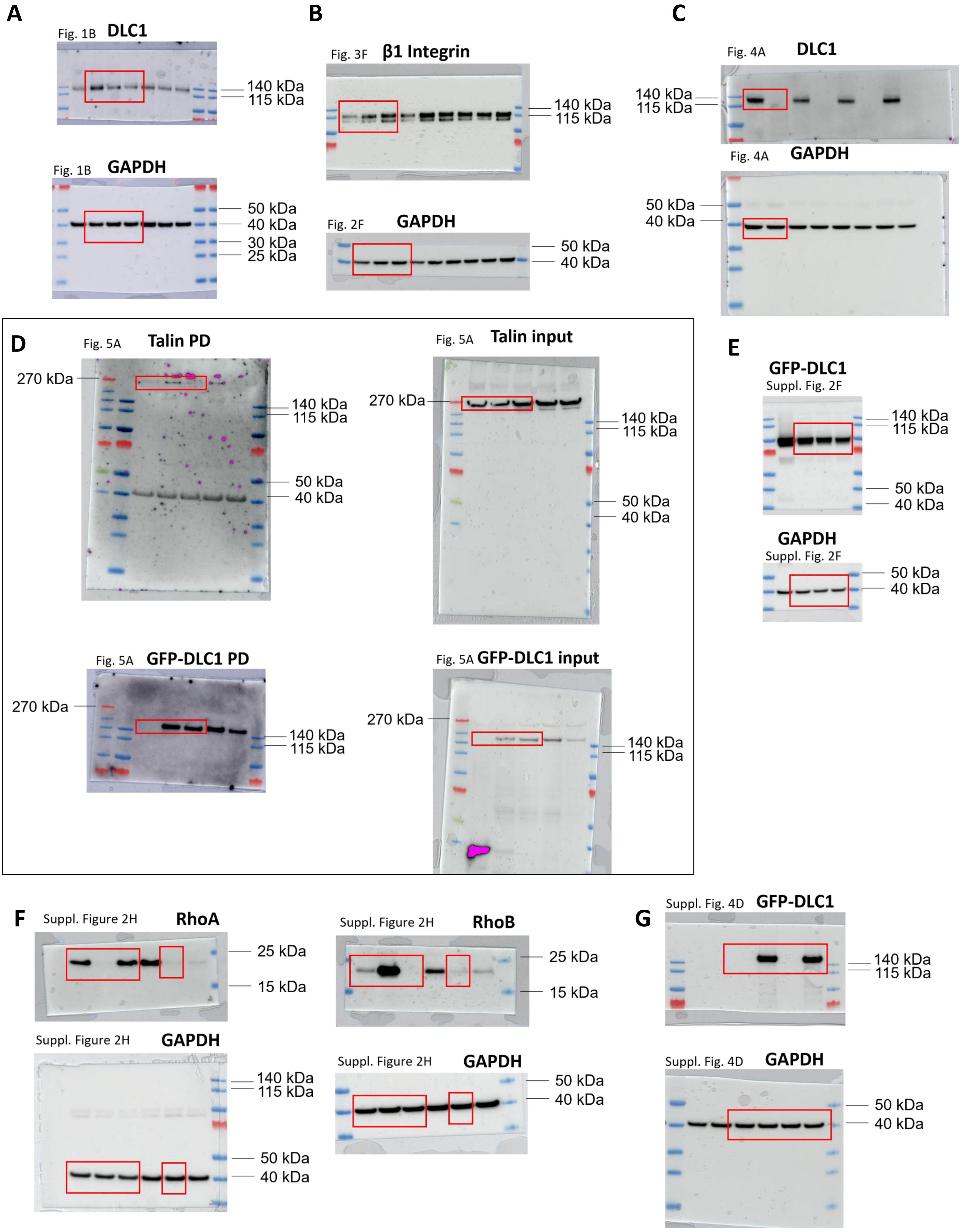


**Supplementary figure 7**

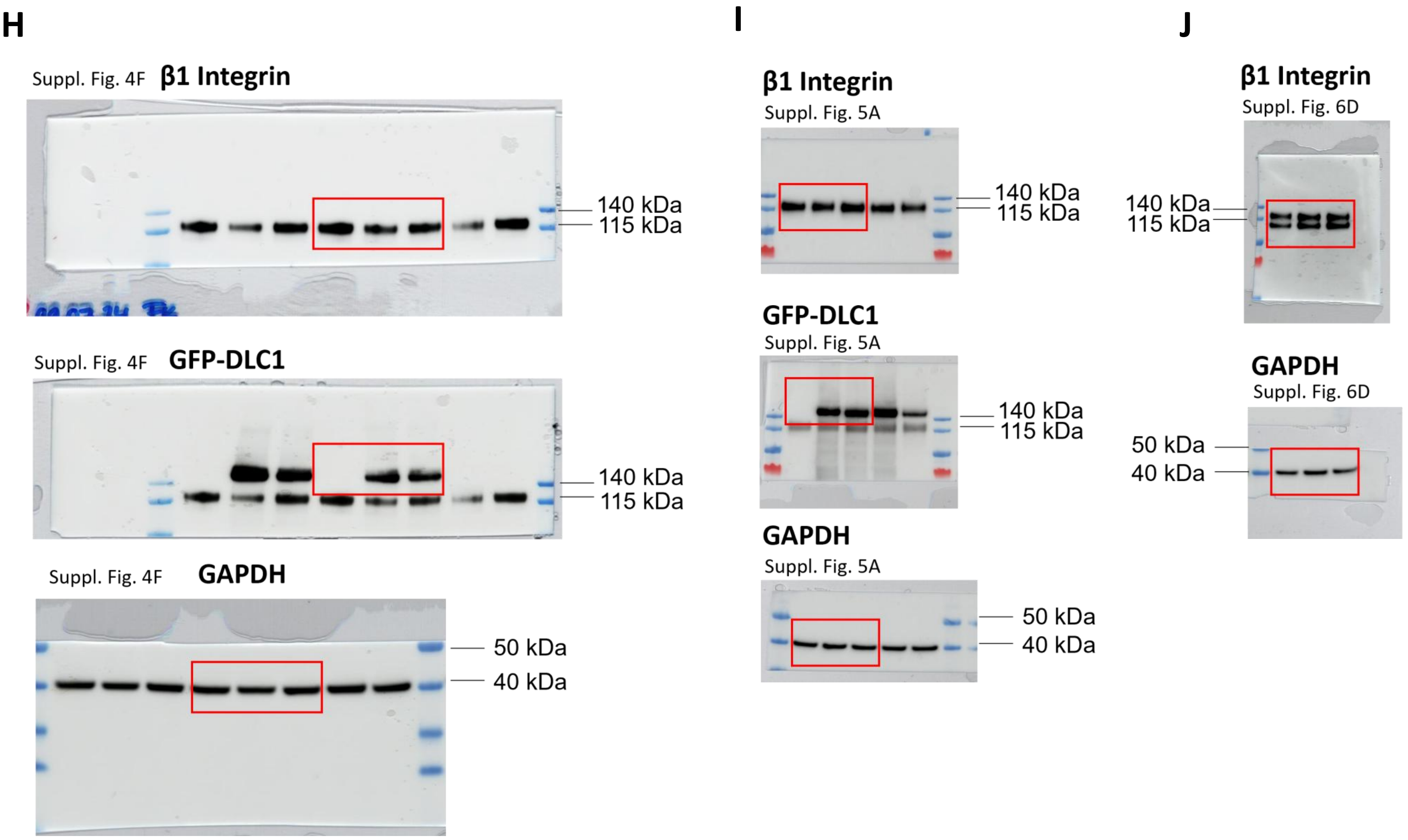


**Supplementary figure 7**
